# Mechanism of Renal Cyst Initiation and Progression Through ETV Transcription Factors and Hedgehog Signaling

**DOI:** 10.64898/2026.08.21.746191

**Authors:** Borum Ryu, Ligyeom Ha, Del L. Dsouza, Erika I. Boesen, Sung-Ho Huh

**Author notes:** Corresponding Author: Sung-Ho Huh, Ph.D., 2500 North State Street, Jackson, MS 39216. Author Contributions: designed experiments (S.H.), performed experiments (S.H., L.H., D.D.), analyzed data (S.H., L.H., D.D., E.B., B.R.), wrote the paper (B.R., E.B., S.H.).

## Abstract

Renal cysts are categorized as non-pathogenic simple cysts and pathogenic malignant cysts based on their pathophysiological status. Cyst formation is divided by cyst initiation and cyst progression/promotion. Pathogenic cysts are thought to be developed through continuous initiation followed by progression until pathogenic status is achieved. Although many genetic and environmental factors are identified to cause pathogenic cyst formation, the mechanisms that discriminate cyst initiation and progression are poorly understood.

Using genetic mutation models of ETV transcription factors, ETV1, ETV4, and ETV5, and a pharmacological inhibitor of hedgehog signaling, cyclopamine, we identified one of the mechanisms regulating cyst initiation and progression. Nephron specific deletion of ETV4 and ETV5 initiated cyst formation. However, cyst initiation did not continue as animals grow, and a limited number of the initial cysts underwent further growth. Additional deletion of ETV1 was required for continuous initiation in addition to promotion of cyst growth. Furthermore, administration of cyclopamine attenuated promotion of cyst progression but had little effect on cyst initiation.

Therefore, we provide evidence that cyst initiation and progression is genetically and molecularly distinct and can be modulated. This information provides new insight into how to control renal cyst initiation and progression and can be used to suppress pathogenic cyst growth.

## Introduction

Renal cysts are characterized by fluid-filled sacs developing in the kidney. Aberrant cyst growth and multi-cyst formation destroys the surrounding renal parenchyma, leading to end-stage kidney disease. Therefore, cystogenesis is the hallmark of the pathogenesis of various kidney diseases including polycystic kidney disease (PKD), glomerulocystic kidney disease GCKD, medullary cystic kidney diseases MCKD, and nephronophthisis [1–3]. Renal cysts develop throughout the lifetime, including developmental stages, and earlier cyst development often results in more severe pathology [4].

ETV1, ETV4, and ETV5 genes are members of the polyomavirus enhancer activator (Pea) 3 subfamily of ETS domain containing transcription factors, which share sequence homology and DNA-binding affinity [5–7]. During murine kidney development, *Etv4* and *Etv5* are highly expressed in ureteric buds (UBs) and identified as functional downstream targets of GDNF/Ret to promote UB induction and elongation [8–10]. However, *Etv4* and *Etv5* are not only expressed in UBs but also expressed in metanephric mesenchyme, which contains nephron progenitor cells (NPCs) [8]. Furthermore, *Etv4* and *Etv5a* are required for multi-ciliated cell generation and differentiation from NPCs during zebrafish kidney development [11]. We identified that FGF20 in human and FGF9 and FGF20 in mouse are required for NPC maintenance and that *Etv4* and *Etv5* are downregulated upon loss of *Fgf9* and *Fgf20* in developing mouse kidneys [12]. However, the roles of ETV transcription factors in metanephric mesenchyme and subsequent nephron compartments in higher vertebrates are not known.

Here, we investigated the role of ETV1, ETV4, and ETV5 in the metanephric mesenchyme and nephron compartments by analyzing metanephric mesenchyme specific *Etv1* and *Etv5* compound mutants together with a conventional deletion mutant of *Etv4*. We identified that ETV transcription factors are dispensable to NPC maintenance. However, we identified that ETV transcription factors suppressed renal cystogenesis originating from nephron compartments. The cysts showed stereotypic cellular features of cystic kidney diseases. Further analyses indicated that loss of *Etv4* and *Etv5* triggers cyst initiation, but a very limited number of cysts undergo further progression. We identified that further loss of *Etv1* is required for continuous cyst initiation and further progression. In addition, we identified individual cyst growth is partly due to activation of Hedgehog (Hh) signaling. Our study suggests that cyst initiation and progression can be regulated independently. This information will be useful to understand to define cyst initiation and progression. Additionally, this information may be used to diagnose cyst progression and/or treat to lower cystic burden in various cystic diseases.

## Results

### *Etv4* and *Etv5* are not required for NPC development

We previously identified that the expression of *Etv4* and *Etv5* was diminished in the kidneys of *Fgf9* and *Fgf20* double mutant (*Fgf9^-/-^; Fgf20^-/-^*) embryos [12, 13]. Since FGF9 and FGF20 are required for NPC maintenance, ETV4 and ETV5 also may be required for development of NPCs. Previous studies indicated that both *Etv4* and *Etv5* were expressed in the UBs and NPCs during kidney development [8]. We confirmed that *Etv4* and *Etv5* were expressed in those tissues with higher expression level in UBs than NPCs at P0 (SupFigure 1). To investigate the function of ETV4 and ETV5 in the NPCs, *Etv5* was conditionally deleted with *Fgf20^Cre^*, NPC specific Cre line [12]. *Etv4* null mutant was used to delete both *Etv4* and *Etv5* (*Etv4^-/-^; Etv5^fl/fl^; Fgf20^Cre/+^*, in short DKO). In DKO kidneys at Postnatal 0 (P0), *Etv4* message was completely diminished and *Etv5* message was not detected in NPCs, but expression remained intact in the UBs (SupFigure 1). Different from FGF9/20 double mutant [13], gross morphological and histologic observation indicate that size of kidneys in control (*Etv4^-/+^; Etv5^fl/+^; Fgf20^Cre^*), ETV4-KO (*Etv4^-/-^; Etv5^fl/+^; Fgf20^Cre^*) and ETV5-KO (*Etv4^-/+^; Etv5^fl/fl^; Fgf20^Cre^*) were comparable without showing any renal aplasia or hypoplasia (Figure 1A and B). In addition, immunostaining of NPC marker, Six2 indicates that NPCs were developed normally at P0 kidneys in all genotypes (SupFigure 2), suggesting that the ETV4 and ETV5 are dispensable for NPC development.

**Figure 1.**
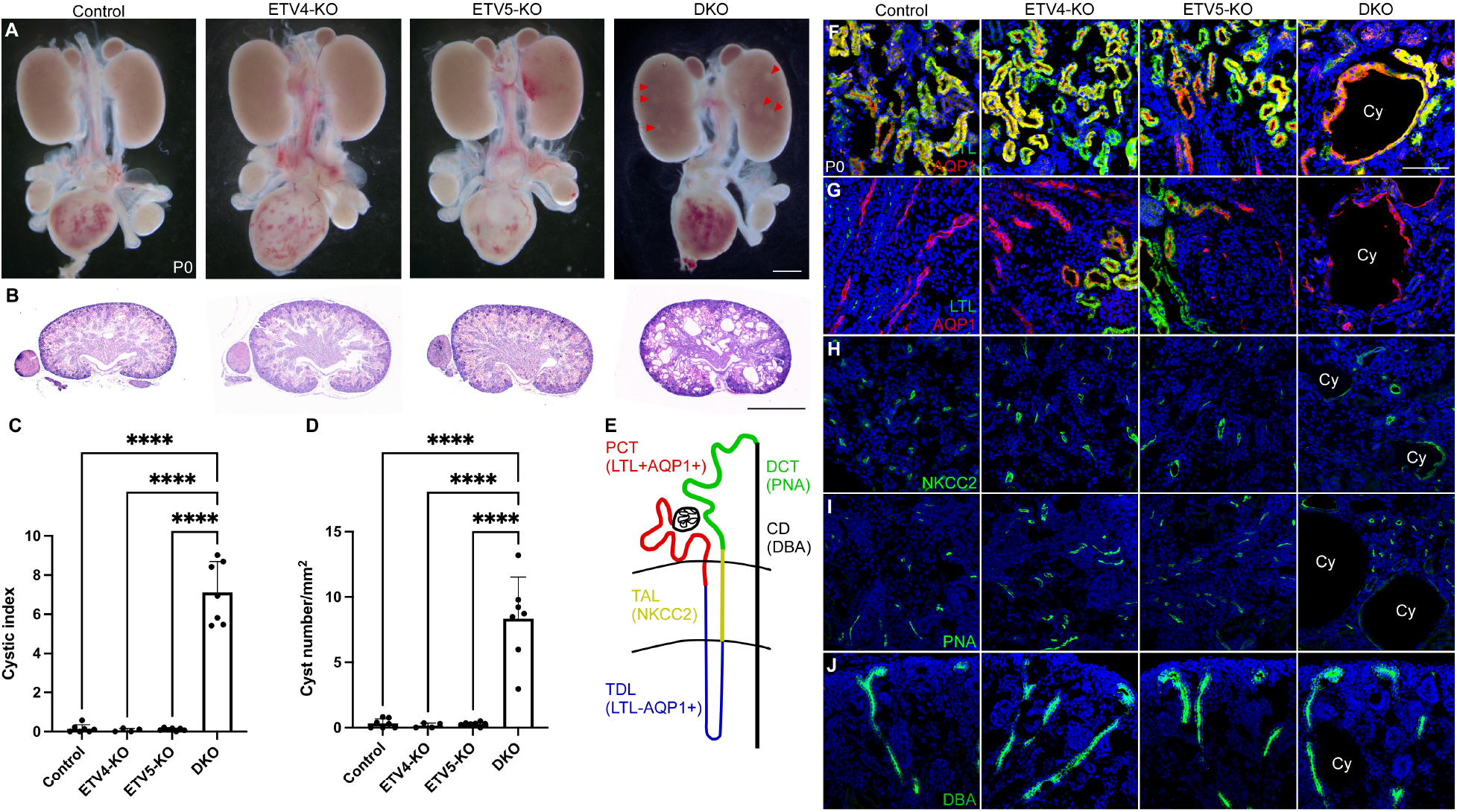
Loss of ETV4 and ETV5 results in renal cystogenesis. (A) Morphology of urogenital system in the newborn *Etv4* and *Etv5* compound mutants. Transparent bubble-like cystic regions (red arrows) are visible in DKO (*Etv4^-/-^; Etv5^fl/fl^; Fgf20^Cre/+^*) compared to grossly undistinguishable morphology of control (*Etv4^-/+^; Etv5^fl/+^; Fgf20^Cre/+^*), ETV4-KO (*Etv4^-/-^; Etv5^fl/+^; Fgf20^Cre/+^*), and ETV5-KO (*Etv4^-/+^; Etv5^fl/fl^; Fgf20^Cre/+^*). (B) Hematoxylin & Eosin staining of the newborn *Etv4* and *Etv5* compound mutants showing polycystic kidneys in DKOs but no other genotypes. Cystic index (C) and cyst number (D) analyses show that DKOs have significantly increased cystic index and cyst number compared to controls, ETV4-KOs, and ETV5-KOs. (E) Schematic diagram of kidney compartments indicating markers used in F-J. (F-J) Immuno-staining of markers indicating specific kidney compartments. LTL and Aquaporin 1 (AQP1) double positive for proximal convoluting tubule (PCT) (F), LTL negative and AQP1 positive for thin descending limb of the loop of Henle (TDL) (G), NKCC2 positive for thick ascending limb of the loop of Henle (TAL) (H), PNA positive for distal convolute tubule (DCT) (I), and DBA for collect duct (CD) (J). Cystic cells (Cy) are labelled with kidney compartment markers of PCT, TAL, TDL, and DCL, indicating they originate from nephron progenitor cells but not CD. ****P < 0.0001. Scale bar, 1mm in A and B, 100µm in F-J.

### Loss of *Etv4* and *Etv5* results in renal cysts

Despite of apparent normal NPC development and kidney size, we observed renal cysts in the kidneys of DKO (Figure 1A and B). Control, ETV4-KO, and ETV5-KO did not show renal cysts (Figure 1A and B). We measured the area and number of cysts (cystic index and cyst number, respectively) using H&E stained kidneys shown in Figure 1B. Cystic index was significantly increased in DKO kidneys compared to all other genotypes (0.15 ± 0.21 in control, n=7; 0.07 ± 0.08 in ETV4-KO, n=4; 0.10 ± 0.07 in ETV5-KO, n=7; and 7.10 ± 1.59 in DKO, n=7, P<0.0001) (Figure 1C). Cyst numbers were also significantly increased in DKO kidneys compared to all other genotypes (0.32 ± 0.35 per mm^2^ in control, n=7, 0.16 ± 0.18 per mm^2^ in ETV4-KO, n=4, 0.27 ± 0.15 per mm^2^ in ETV5-KO, n=7, 8.33 ± 3.18 per mm^2^ in DKO, n=7, P<0.0001) (Figure 1D). Marker study with cell type specific antibodies, lectin, and agglutinins showed cysts originated from nephron compartments including proximal convoluted tubule (PCT) (Figure 1F), thin descending limb of the loop of Henle (TDL) (Figure 1G), thick ascending limb of the loop of Henle (TAL) (Figure 1H), and distal convoluted tubule (DCT) (Figure 1I). Cysts did not develop in collecting duct (Figure 1J). Together, these data indicate that loss of *Etv4* and *Etv5* in NPCs results in cyst formation in the nephron compartment but not in the collecting duct system, which originates from the UB.

### Increased proliferation, cell death, and cilia length in cystic kidneys

Next, we evaluated cellular changes in cystic kidneys by analyzing cell proliferation, cell death, and cilia length. Proliferation rates of the non-cystic kidneys (controls, ETV4-KO, ETV5-KO) were comparable, as determined by Ki67 labeling. However, cystic kidneys of DKO showed a significant increase in proliferation compared to all other genotypes (10.39 ± 1.08 in control, n=4, 12.59 ± 6.0 in ETV4-KO, n=3, 11.14 ± 3.9 in ETV5-KO, n=4, 28.60 ± 5.35 in DKO, n=5, P=0.0078 to control, P=0.0354 to ETV4-KO, P=0.0111 to ETV5-KO) (Figure 2A and B). Cell death, determined by labeling with activated-Caspase3, was also significantly increased in DKO kidneys compared to other genotypes (0.08 ± 0.05 in control, n=5, 0.10 ± 0.02 in ETV4-KO, n=2, 0.06 ± 0.04 in ETV5-KO, n=4, 0.74 ± 0.17 in DKO, n=4, P<0.0001) (Figure 2C and D). Abnormal cilia is a key feature of renal cystogenesis [2, 14]. Therefore, we investigated the length of the cilia labeling with Arl13b, which labels cilia [15, 16]. Cilia length was significantly increased in the cystic kidneys of DKO compared to all other genotypes (4.09 ± 2.46 μm in control, n=783 combined with 4 animals, 4.19 ± 2.17 μm in ETV4-KO, n=793 combined with 3 animals, 4.01 ± 2.13 μm in ETV5-KO, n=879 combined with 4 animals, 5.53 ± 2.88 μm in DKO, n=692 combined with 4 animals, P<0.0001) (Figure 2E and F). Together, these data suggest that loss of ETV4 and ETV5 in the nephron compartment produces cystic disease-like cellular phenotypes.

**Figure 2.**
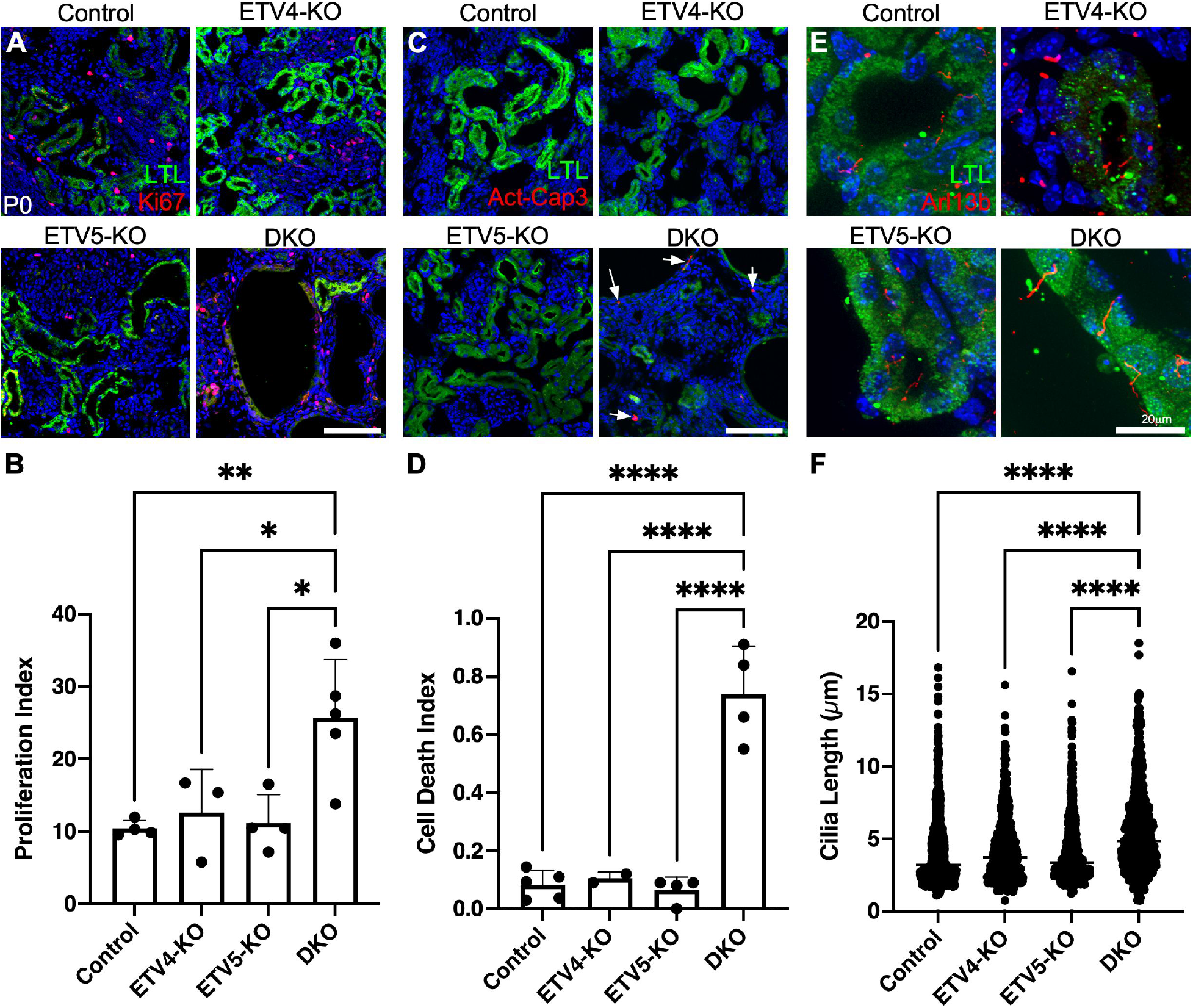
Cystic kidneys increase cell proliferation, cell death, and cilia length in P0 ETV4/5 compound animals. Immunostaining of controls, ETV4-KO, ETV5-KO, and DKO kidneys with LTL and Ki67 for cell proliferation (A), LTL and Activated-Caspase 3 for cell death (Act-Casp3) (C), and LTL and Arl13b (E). Quantification of proliferation index (B), Cell death index (D), and cilia length (F) show increases in DKOs but no other genotypes. **P < 0.005, ****P < 0.0001. Scale bar, 100µm in A and C, 20µm in E.

### Renal cysts in DKO mice regress as animals grow

Next, we investigated whether renal cysts in DKO further progress as the animals grow. Cystic index and cyst number were still significantly increased in the kidneys of DKO at P30 (cystic index, 0.04 ± 0.06 in control, n=5, 0.35 ± 0.32 in ETV4-KO, n=5, 0.40 ± 0.18 in ETV5-KO, n=8, 7.47 ± 6.61 in DKO, n=7, P=0.0087 to control, P=0.0121 to ETV4-KO, P=0.0045 to ETV5-KO; cyst number, 0.11 ± 0.15 per mm^2^ in control, n=5, 0.27± 0.21 per mm^2^ in ETV4-KO, n=5, 0.38 ± 0.19 per mm^2^ in ETV5-KO, n=8, 1.5 ± 1.04 per mm^2^ in DKO, n=7, P=0.0028 to control, P=0.0082 to ETV4-KO, P=0.0084 to ETV5-KO) and at P180 animals (Cystic index, 0.14 ± 0.16 in control, n=17, 0.22 ± 0.19 in ETV4-KO, n=5, 0.28 ± 0.19 in ETV5-KO, n=9, 1.07 ± 1.34 in DKO, n=11, P=0.0079 to control, P=0.1325 to ETV4-KO, P=0.0771 to ETV5-KO; cyst number, 0.07 ± 0.06 per mm^2^ in control, n=17, 0.13± 0.20 per mm^2^ in ETV4-KO, n=5, 0.17 ± 0.13 per mm^2^ in ETV5-KO, n=9, 0.37 ± 0.19 per mm^2^ in DKO, n=11, P<0.0001 to control, P=0.0052 to ETV4-KO, P=0.0060 to ETV5-KO) compared to control, ETV4-KO, and ETV5-KO (Figure 3A-F). However, when we compared kidneys of DKO among P0, P30, and P180, we observed a gradual decrease in cystic index and cyst number as animals aged. Cystic index was comparable between P0 and P30 animals but significantly decreased in P180 animals compared to both P0 and P30 animals (7.10 ± 1.59 in P0, n=7, 7.47 ± 6.61 in P30, n=7, 1.07 ± 1.34 in P180, n=11, P=0.0068 to P0, P=0.0042 to P30) (Figure 3G). In addition, cyst number was significantly decreased in P30 animals compared to P0 animals (8.33 ± 3.18 per mm^2^ in P0, n=7, 1.5 ± 1.04 per mm^2^ in P30, n=7, P<0.0001), and further decreased in P180 animals compared to P0 and P30 animals (8.33 ± 3.18 per mm^2^ in P0, n=7, 1.5 ± 1.04 per mm^2^ in P30, n=7, 0.37 ± 0.19 per mm^2^ in P180, n=11, P<0.0001 to P0 and P30, respectively) (Figure 3H). We still observed the significant increase of cell proliferation, cell death, and cilia length in P30 DKO kidneys compared to control, ETV4-KO and ETV5-KO (cell proliferation, 3.66 ± 1.58 in control, n=5, 3.55 ± 3.23 in ETV4-KO, n=3, 5.27 ± 2.91 in ETV5-KO, n=4, 10.37 ± 1.25 in DKO, n=3, P=0.0103; cell death, 0.02 ± 0.02 in control, n=5, 0.04 ± 0.03 in ETV4-KO, n=3, 0.02 ± 0.02 in ETV5-KO, n=4, 0.53 ± 0.23 in DKO, n=3, P<0.0001; cilia length, 3.78 ± 1.37 μm in control, n=927 combined with 5 animals, 3.84 ± 1.37 μm in ETV4-KO, n=369 combined with 3 animals, P<0.0001, 3.91 ± 1.52 μm in ETV5-KO, n=823 combined with 4 animals, P<0.0001, 5.91 ± 3.30 μm in DKO, n=439 combined with 4 animals, P<0.0001 to control) (SupFigure 3). These data indicate that many cysts do not progress in DKO animals, although the growing cysts still show stereotypic cellular phenotypes.

**Figure 3.**
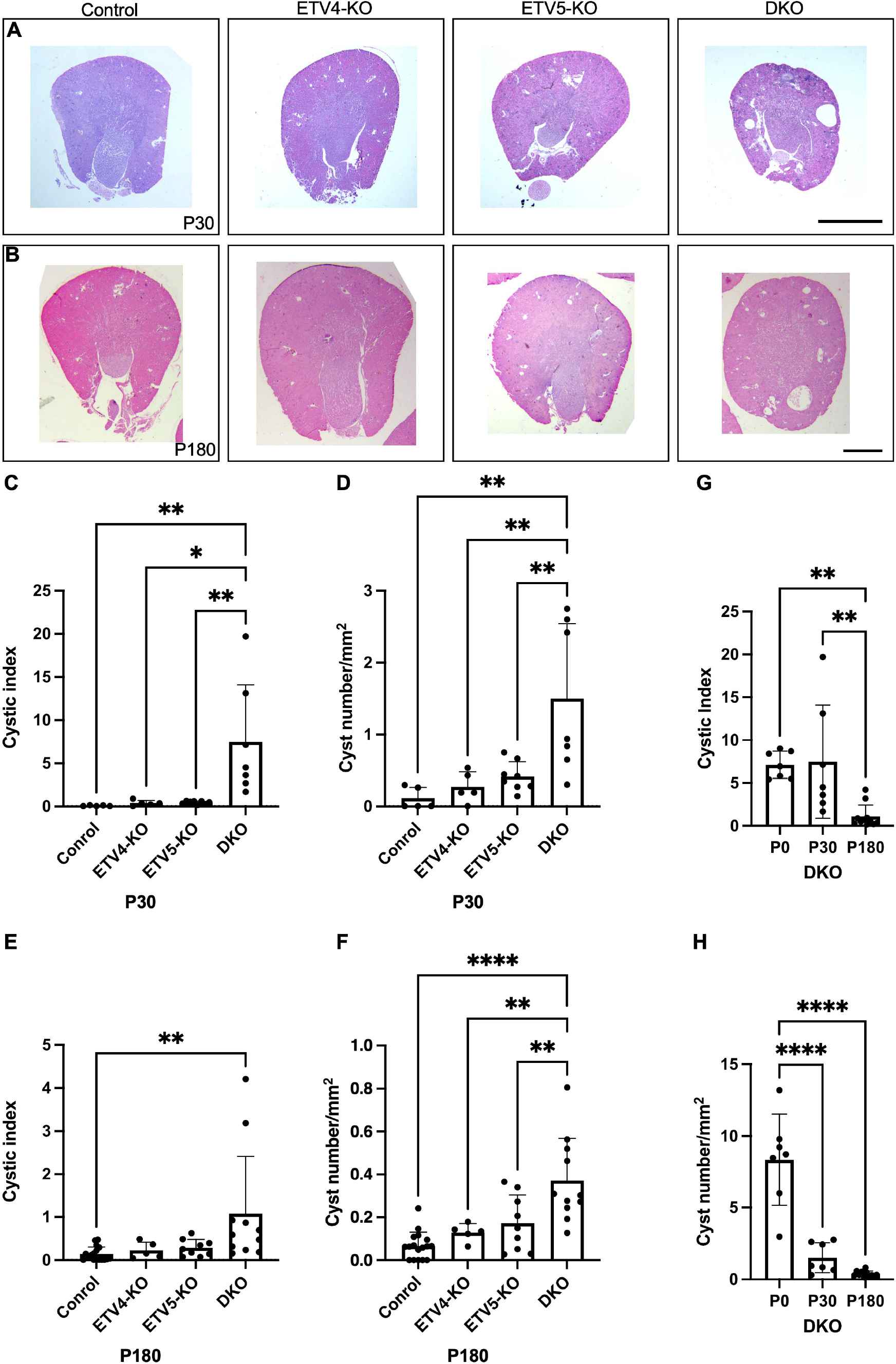
Cyst sizes are increased but number of cysts is decreased as DKOs grow. (A and B) Hematoxylin & Eosin staining of the P30 (A) and P180 (B) controls, ETV4-KO, ETV5-KO, and DKO kidneys showing cysts in DKO kidneys. No cysts are shown in controls, ETV4-KO, and ETV5-KO kidneys. (C-F) Measuring cystic index and cyst number of P30 and P180 kidneys indicate that DKO kidneys show increased cystic index and cyst number compared to controls, ETV4-KOs, and ETV5-KOs. (G and H) comparison of cystic index (G) and cyst number (H) among P0, P30, and P30 DKO kidneys showing that gradual decrease in cystic index and cyst number as mice age. *P < 0.05, **P < 0.005, P < 0.0005, ****P < 0.0001. Scale bar, 1mm

### Cell identity is diminished in cysts in DKOs as animal grow

Different from P0 animals, cystic cells of P30 kidneys were only positive for DCT marker, PNA (SupFigure 4D) and negative for all other nephron segment markers including LTL (SupFigure 4A), Aquaporin1 (SupFigure 4B), NKCC2 (SupFigure 4C), in addition to the collecting duct marker DBA (SupFigure 4E). Therefore, we investigated the origin of cysts as animals grow. To investigate whether cystic cells originated from NPCs, we traced the lineage of cystic cells using Rosa-TdTomato mouse line along with *FGF20^Cre^*. At P0, all cystic cells were labelled with tdTomato, and these tdTomato positive cells were detected in NPCs and nephron compartments (labeled with Aquaporin 1, LTL, NKCC2, and Six2) (SupFigure 5A-D). However, tdTomato-positive cells were not detected in the collecting duct cells (labeled with DBA and Aquaporin 2), the endothelial cells (labeled with CD34), nor the fibroblast progenitor cells (labeled with Foxd1) (SupFigure 5E-H). These data indicate that cystic cells originated from NPCs or nephron compartments. At P30, as with P0, all nephron compartments are were labeled with TdTomato in the kidneys of the control mice (SupFigure 6A). All cystic cells in DKO kidneys were also labeled with TdTomato (SupFigure 6B) but lacked staining for nephron segment markers. These data indicate that the cystic cells originated from the nephron compartment and this origin has not been changed at P30.

### ETV1 compensates for ETV4 and ETV5 beyond postnatal age

We questioned whether renal cysts in DKO animals regressed due to compensation from other molecule to protect against the loss of ETV4 and ETV5. ETV1 is a transcription factor closely related to ETV4 and ETV5 in sequence homology and shares the common DNA consensus sequence [17]. In addition, ETV1 cooperates with ETV4 and ETV5 in the development of various organs and cancer [7, 18, 19].Therefore, it is possible that ETV1 may compensate for the loss of ETV4 and ETV5 in kidney development. First, we investigated whether ETV1 is ectopically expressed in the kidneys upon loss of ETV4 and ETV5. mRNA *in situ* hybridization analysis showed that *Etv1* was sparsely expressed in the P0 kidneys of control mice (Figure 4A). Using fluorescence *Etv1* mRNA *in situ* hybridization, we observed ectopic expression of *Etv1* in cystic cells from the DKO kidneys (Figure 4A). In addition, real-time PCR analysis showed that the expression of *Etv1* was significantly increased in the P30 DKO kidneys compared to controls (1.13 ± 0.60 in control, n=6, 2.68 ± 1.13 in DKO, n=6, P=0.0139) (Figure 4B). These data indicate that ETV1 is ectopically induced upon loss of ETV4 and ETV5.

**Figure 4.**
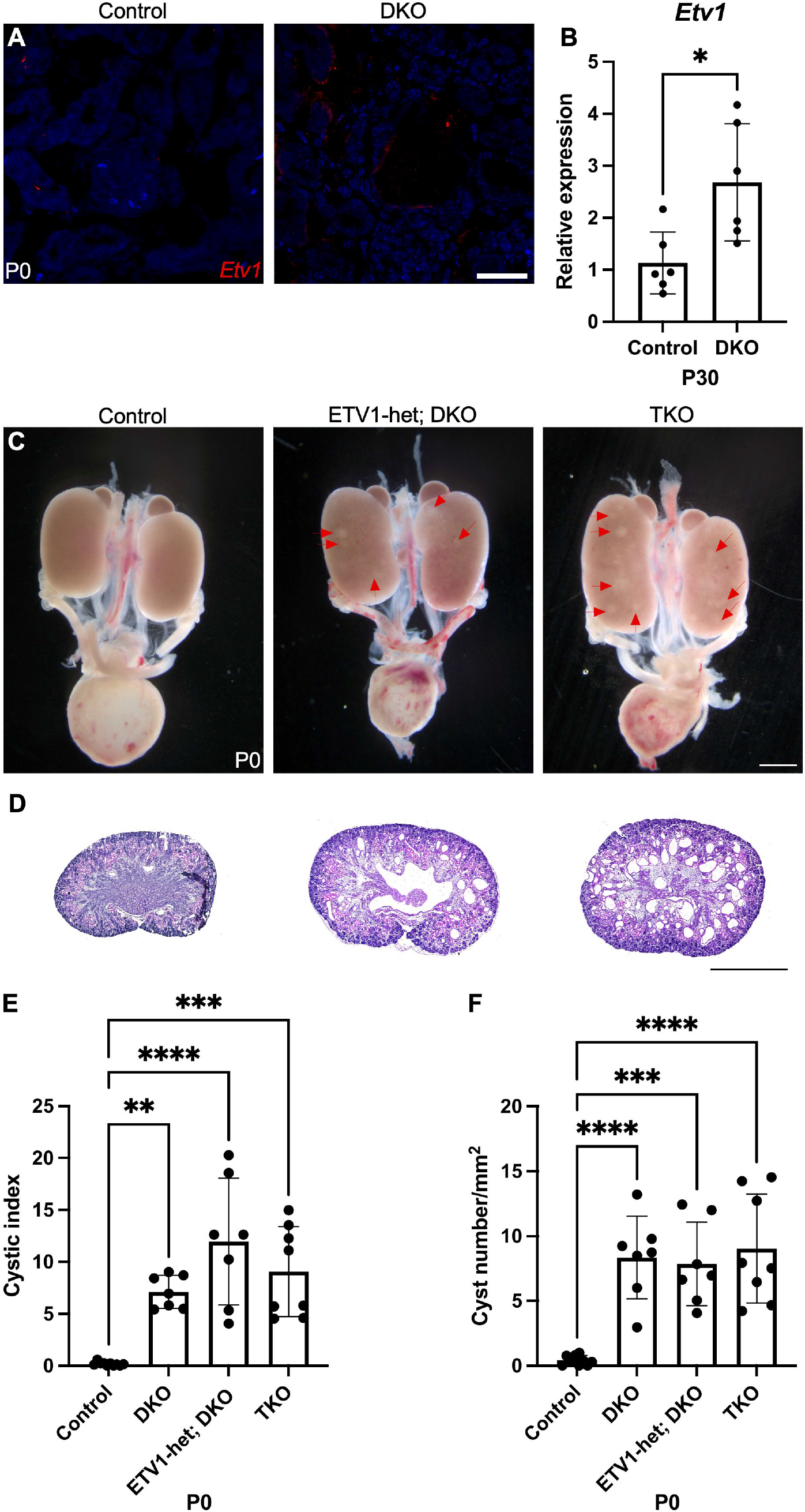
*Etv1* is dispensable during initiation of cystogenesis caused by loss of *Etv4* and *Etv5*. (A) Fluorescent *in situ* hybridization of *Etv1* showing that *Etv1* mRNA is detectable in cystic cells. (B) Quantitative Real-Time PCR of P30 kidneys showing that expression of *Etv1* is increased in DKOs compared to controls. (C) Morphology of urogenital system in the newborn *Etv1*, *Etv4*, and *Etv5* compound mutants. Transparent bubble-like cystic regions (red arrows) are visible in ETV1-het; DKO (*Etv1^fl/+^; Etv4^-/-^; Etv5^fl/fl^; Fgf20^Cre/+^*) and TKO (*Etv1^fl/fl^; Etv4^-/-^; Etv5^fl/fl^; Fgf20^Cre/+^*) compared to control (*Etv1^fl/+^; Etv4^-/+^; Etv5^fl/+^; Fgf20^Cre/+^*). (D) Hematoxylin & eosin staining of the newborn *Etv1*, *Etv4*, and *Etv5* compound mutants showing polycystic kidneys in ETV1-het; DKOs and TKOs. Comparison of cystic index (E) and cyst number (F) of controls, DKO, ETV1-Het; DKO, and TKO kidneys showing that cystic index and cyst number are comparable among DKO, ETV1-het; DKO, and TKO and increased compared to controls. *P < 0.05, **P < 0.005, P < 0.0005, ****P < 0.0001. Scale bar, 100µm in A and C, 1mm in C and D.

Next, we questioned whether additional loss of ETV1 in NPCs exacerbates the defects of kidneys in DKOs. Using an *Etv1^fl^* mouse line, we conditionally deleted either one copy or two copies of *Etv1* in the NPC population under DKO conditions. Deleting all three ETVs (*Etv1^fl/fl^; Etv4^-/-^; Etv5^fl;fl^; Fgf20^Cre^*, TKO) did not show significant NPC population changes at P0 (SupFigure 7), indicating that all three ETVs are dispensable to generate and maintain NPCs. Gross morphology and histological analysis of P0 animals showed that both ETV1het; DKO (*Etv1^fl/+^; Etv4^-/-^; Etv5^fl;fl^; Fgf20^Cre^*) and TKO exhibited multiple cysts in both kidneys (Figure 4C and D). Cystic index and cyst number of DKO, ETV1het; DKO, and TKO were comparable to each other, but significantly increased compared to controls (cystic index, 0.16 ± 0.17 in control, n=10, 7.10 ± 1.59 in DKO, n=7, P=0.0031, 11.95 ± 6.10 in ETV1het; DKO, n=7, P<0.0001, 9.06 ± 4.34 in TKO, n=8, P=0.0001; cyst number, 0.42 ± 0.37 per mm^2^ in control, n=10, 8.33 ± 3.19 per mm^2^ in DKO, n=7, P<0.0001, 10.83 ± 6.83 per mm^2^ in ETV1het; DKO, n=7, P=0.0001, 9.02 ± 4.20 per mm^2^ in TKO, n=8, P<0.0001) (Figure 4E and F). Proliferation rate was significantly increased in ETV1-het; DKO and TKO kidneys compared to control (23.23 ± 9.19 in control, n=4, 42.71 ± 6.79 in ETV1het; DKO, n=5, P=0.0207, 43.24 ± 10.72 in TKO, n=6, P=0.0147) and comparable between ETV1het; DKO and TKO (SupFigure 8A and 8B). Cell death was also significantly increased in ETV1-het; DKO and TKO kidneys compared to controls (0.07 ± 0.05 in control, n=4, 0.59 ± 0.17 in ETV1het; DKO, n=5, P=0.0032, 0.83 ± 0.48 in TKO, n=7, P=0.0161). Cell death of TKO kidneys was comparable to ETV1-het; DKO (SupFigure 8C and 8D). Cilia length was significantly increased in ETV1-het; DKO and TKO kidneys compared to controls (3.84 ± 2.05 μm in control, n=940 combined with 4 animals, 5.24 ± 3.24 μm in ETV1het; DKO, n=683 combined with 5 animals, P<0.0001, 5.75 ± 3.37 μm in TKO, n=1449 combined with 7 animals, P<0.0001). Cilia length of TKO kidney was significantly increased compared to ETV1-het; DKO (P=0.0006) (SupFigure 8E and 8F). Together these data indicate that ETV1 does not influence prenatal renal cystogenesis caused by loss of ETV4 and ETV5.

However, gross morphology and histology showed that by P30, ETV1-het; DKO and TKO kidneys were more transparent and highly cystic compared to control and DKO (Figure 5A and B). Cystic index was significantly increased in both ETV1-het; DKO and TKO kidneys compared to control and DKO (cystic index, 0.04 ± 0.06 in control, n=14, 6.19 ± 4.20 in DKO, n=5, 20.85 ± 12.01 in ETV1het; DKO, n=8, P<0.0001, 25.75 ± 16.34 in TKO, n=6, P<0.0001) and compared to DKO (in ETV1-het; DKO, P=0.045, in TKO, P=0.0071) (Figure 5C). Cyst number was also significantly increased in both ETV1-het; DKO and TKO kidneys compared to control (0.03 ± 0.06 per mm^2^ in control, n=12, 1.70 ± 0.98 per mm^2^ in DKO, n=6, 4.32 ± 1.58 per mm^2^ in ETV1het; DKO, n=10, P<0.0001, 4.79 ± 2.86 per mm^2^ in TKO, n=8, P<0.0001) and compared to DKO (in ETV1-het; DKO, P=0.0186, in TKO, P=0.0067) (Figure 5D). Interestingly, comparison of cyst size of DKO, ETV1-het; DKO, and TKO showed that they were comparable to each other (47985 ± 117774 μm^2^, n=104 in 6 DKO animals, 39276 ± 129800 μm^2^, n=816 in 10 ETV1het; DKO animals, 48878 ± 139400 μm^2^, n=543 in 8 TKO animals) (Figure 5E). These data indicate that increased cystic index in ETV1-het; DKO and TKO compared to DKO kidneys are mainly due to increased cyst initiation rather than cyst progression.

**Figure 5.**
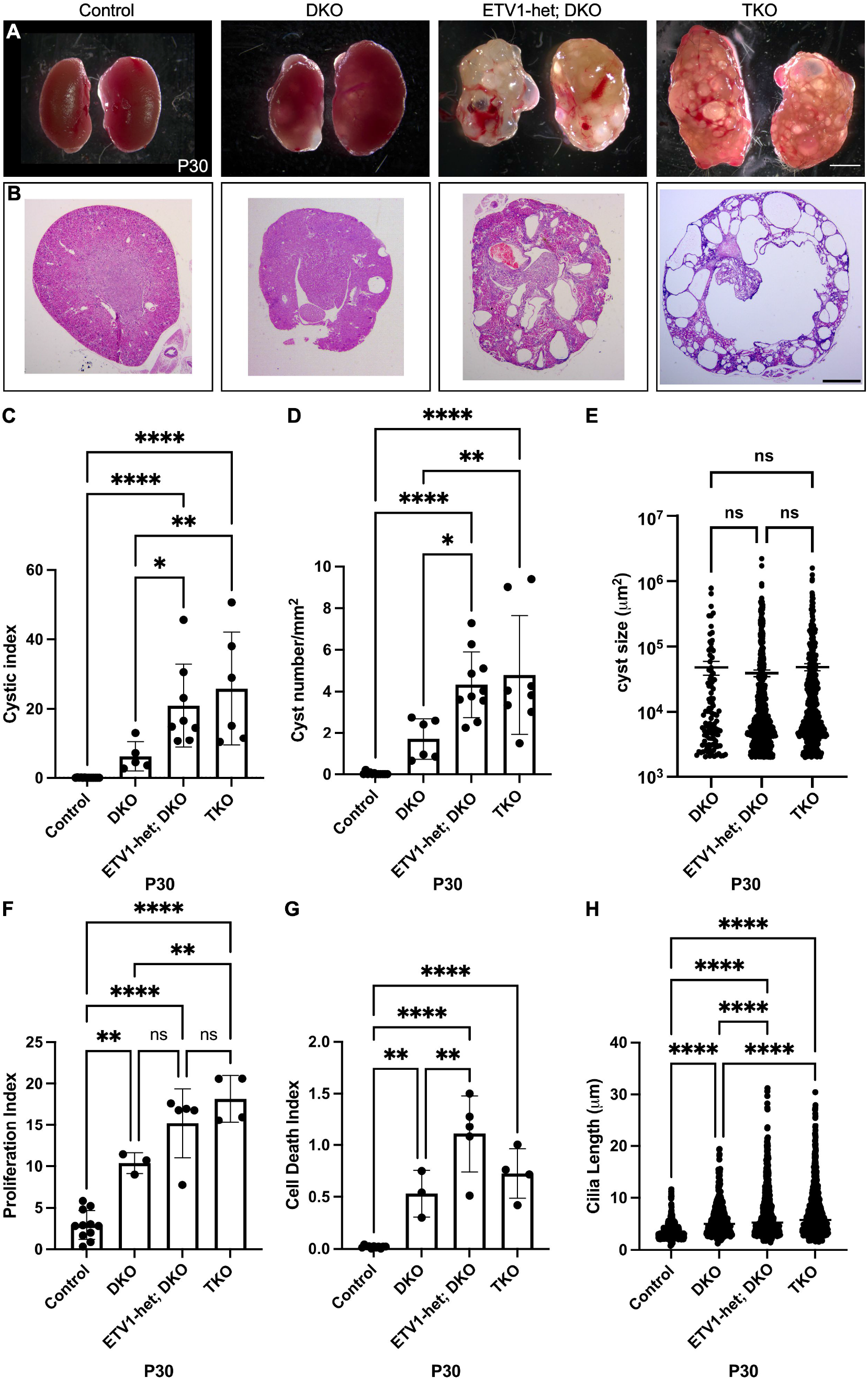
Deletion of ETV1 facilitates cyst progression. (A and B) Gross morphology (A) and hematoxylin & eosin staining (B) of kidneys of P30 control, DKO, ETV1-het; DKO, and TKO animals showing more severe renal cyst formation in ETV1-het; DKO and TKO compared to DKO. (C-G) measurement of cystic index (C), cyst number (D), proliferation index (E), cell death index (F), and cilia length (G) of P30 control, DKO, ETV1-het; DKO, and TKO kidneys showing increase in all categories in ETV1-het; DKO and TKO compared to DKO. *P < 0.05, **P < 0.005, P < 0.0005, ****P < 0.0001. Scale bar, 1mm

Cellular phenotypes were also analyzed at P30. Proliferation rate was significantly increased in DKO, ETV1-het; DKO and TKO kidneys compared to control (2.92 ± 1.74 in control, n=11, 10.37 ± 1.25 in DKO, n=3, P=0.0015, 15.18 ± 4.18 in ETV1het; DKO, n=5, P<0.0001, 18.17 ± 2.80 in TKO, n=4, P<0.0001) and in TKO compared to DKO (P=0.0043) (Figure 5F, SupFigure 9A and B). Cell death was increased in DKO, ETV1-het; DKO and TKO kidneys compared to control (0.02 ± 0.01 in control, n=11, 0.53 ± 0.22 in DKO, n=3, P=0.0062, 1.11 ± 0.37 in ETV1het; DKO, n=5, P<0.0001 to control, P=0.0059 to DKO, 0.72 ± 0.24 in TKO, n=4, P<0.0001), and was increased in ETV1-het; DKO compared to DKO (P=0.0020), and was increased in in ETV1-het; DKO and TKO kidneys compared to control (0.02 ± 0.01 in control, n=6, 1.11 ± 0.37 in ETV1-het; DKO, n=5, P<0.0001, 0.72 ± 0.24 in TKO, n=4, P=0.0020) (Figure 5G, SupFigure 9C and D). Cilia length was increased in DKO, ETV1-het; DKO, and TKO kidneys compared to control (3.51 ± 1.42 μm in control, n=767 combined with 5 animals, 5.91 ± 3.30 μm in DKO, n=439 combined with 4 animals, P<0.0001 to control, ETV1-het; DKO, and TKO, 7.12 ± 4.97 μm in ETV1-het; DKO, n=645 combined with 6 animals, P<0.0001 to control and DKO, 7.10 ± 4.58 μm in TKO, n=842 combined with 6 animals, P< 0.0001 to control and DKO), increased in ETV1-het; DKO and TKO compared to DKO (P<0.0001 to all) (Figure 5H, SupFigure 9E and F). Together these data shows that deletion of ETV1 enhances cellular defects leading to renal cysts.

We also examined identities of cystic cells of ETV1-het; DKO and TKO kidneys. Different from cystic cells in DKO kidneys which only labeled with marker of DCT (PNA), cystic cells in ETV1-het; DKO and TKO kidneys were labeled with all nephron compartments including LTL, NKCC2, and PNA but not collecting duct marker, DBA (SupFigure 10). These data suggest that maintaining cell identity may be required for cyst initiation.

### Loss of ETVs causes fatal chronic kidney disease

Based on the renal phenotypes observed, which resemble pathogenic cystic kidney diseases, we investigated whether ETV mutant mice exhibit renal pathogenic symptoms. First, we determined survival rate of the ETV compound mutants. No control and DKO animals died prior to 6 months (Figure 6A). However, we noticed 50% of ETV1-het; DKO and 30% of TKO animals showed early mortality, beginning at 1 month and 3 months, respectively (Figure 6A). Gene ontology analyses of transcriptomes of P30 control and TKO kidneys showed downregulation of mitochondrial function (ex, oxidative phosphorylation), and ion homeostasis, transport, and excretion (ex, inorganic ion transmembrane transport, proton transmembrane transport, transport of small molecule, and cation homeostasis) in TKO kidneys compared to controls (Figure 6B). On the other hand, DNA synthesis, DNA damage maintenance, cell division, microtubule and centriole assembly were increased in TKO kidneys compared to controls (Figure 6C).

**Figure 6.**
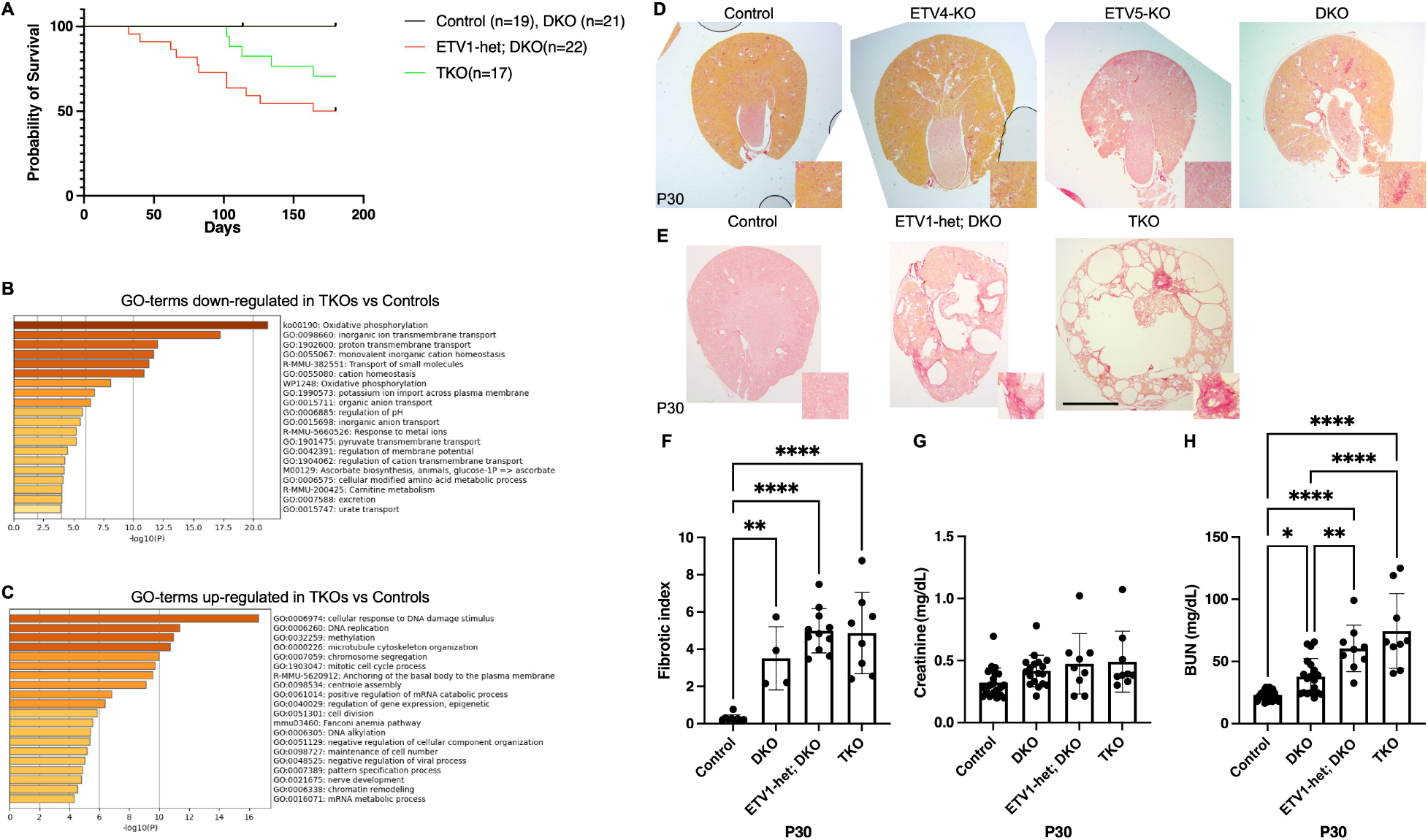
Loss of ETVs produces severe renal pathology. (A) Survival curve of controls, DKOs, ETV1-het; DKOs, and TKOs showing decreased survival in ETV1-het; DKO and TKO animals. (B and C) Gene ontology (GO) analyses of transcriptome of P30 controls and TKOs showing decreased kidney function and increased cell damage. (D and E) Sirius Red staining showing fibrosis in P30 DKO, ETV1-het; DKO, and TKO kidneys. (F-H) analyses of fibrotic index (F), creatinine concentration (G), and BUN (H) of P30 control, DKO, ETV1-het; DKO, and TKO kidneys. *P < 0.05, **P < 0.005, P < 0.0005, ****P < 0.0001. Scale bar, 1mm

In addition, Sirius Red staining showed that kidneys of DKO, ETV1-het; DKO, and TKO animals were fibrotic (Figure 6D and E). Fibrotic index (by measuring Sirius Red positive area) was increased in DKO, ETV1-het; DKO, and TKO kidneys compared to control (0.29 ± 0.18 in control, n=9, 3.50 ± 1.70 in DKO, n=4, P=0.004, 4.98 ± 1.18 in ETV1-het; DKO, n=11, P<0.0001, 4.83 ± 2.18 in TKO, n=8, P<0.0001) (Figure 6F). Creatinine concentration was variable but not statistically different among all genotypes (0.33 ± 0.13 mg/dL in control, n=18, 0.415 ± 0.127 mg/dL in DKO, n=18, 0.472 ± 0.243 mg/dL in ETV1-het; DKO, n=9, 0.490 ± 0.246 mg/dL in TKO, n=9) (Figure 6G). However, blood urea nitrogen (BUN) level was significantly increased in ETV1-het; DKO and TKO animals compared to control and DKO (22.49 ± 3.83 mg/dL in control, n=19, 37.56 ± 14.58 mg/dL in DKO, n=19, P=0.0334 to control, 60.34 ± 18.69 mg/dL in ETV1-het; DKO, n=9, P<0.0001 to control and P=0.0066 to DKO, 74.37 ± 30.11 mg/dL in TKO, n=9, P<0.0001 to control and DKO), in addition to being increased in DKO mice compared to control (Figure 6H). Together, these data indicate that ETV compound mutants suffer chronic kidney disease.

### Activation of hedgehog signaling is required for cyst elongation progression

In some polycystic kidney models, ectopic activation of hedgehog signaling promotes cyst growth [20, 21]. Therefore, we investigated whether hedgehog signaling was induced in ETV compound mutants. Real-time PCR analyses of hedgehog signal-related genes showed that the expression of *Shh* was significantly increased in TKO kidneys compared to control and ETV1-het; DKO (1.03 ± 0.27 in control, n=4, 1.50 ± 0.81 in ETV1-het; DKO, n=5, 2.42 ± 0.27 in TKO, n=5, P=0.0066 to control and P=0.0491 to ETV1-het; DKO), the expression of *Gli2* was significantly increased in TKO kidneys compared to control (1.01 ± 0.31 in control, n=4, 1.47 ± 0.53 in ETV1het; DKO, n=5, 1.81 ± 0.31 in TKO, n=5, P=0.0234), the expression of *Ptch1* was significantly decreased in ETV1-het; DKO kidneys compared to control (1.01 ± 0.19 in control, n=4, 0.54 ± 0.24 in ETV1-het; DKO, n=5, P=0.0062, 0.71 ± 0.08 in TKO, n=5), and the expression of *Ptch2* was significantly decreased in ETV1-het; DKO and TKO kidneys compared to control (1.00 ± 0.08 in control, n=4, 0.59 ± 0.21 in ETV1-het; DKO, n=5, P=0.0093, 0.64 ± 0.33 in TKO, n=5, P=0.0036) at P0 (SupFigure 11). Expressions of *Smo*, *Ptch2*, and *Gli3* were comparable among genotypes (*Smo*, 1.01 ± 0.15 in control, n=4, 1.18 ± 0.23 in ETV1-het; DKO, n=5, 0.84 ± 0.25 in TKO, n=5, *Gli3*,1.01 ± 0.17 in control, n=4, 1.21 ± 0.38 in ETV1-het; DKO, n=5, 1.17 ± 0.08 in TKO, n=4). At P30, expressions of *Shh*, *Ptch2*, *Gli1*, *Gli2*, and *Gli3* were significantly increased in ETV1-het; DKO and TKO kidneys compared to controls (*Shh*, 0.74 ± 0.50 in control, n=8, 2.92 ± 1.52 in ETV1-het; DKO, n=8, P=0.0015, 2.15 ± 0.94 in TKO, n=8, P=0.0391; *Ptch2*, 0.93 ± 0.36 in control, n=8, 3.64 ± 1.84 in ETV1-het; DKO, n=7, P=0.0005, 3.63 ± 1.32 in TKO, n=7, P=0.0029; *Gli1*, 0.91 ± 0.27 in control, n=8, 2.70 ± 1.04 in ETV1-het; DKO, n=8, P=0.0002, 1.91 ± 0.62 in TKO, n=8, P=0.0280; *Gli2*, 1.03 ± 0.27 in control, n=6, 4.34 ± 1.01 in ETV1-het; DKO, n=4, P=0.0027, 3.68 ± 1.95 in TKO, n=4, P=0.0119; *Gli3*, 1.04 ± 0.34 in control, n=8, 7.04 ± 3.97 in ETV1-het; DKO, n=7, P=0.0417, 10.30 ± 7.52 in TKO, n=6, P=0.0045) (Figure 7A, D, E, F, G). Expressions of *Smo* and *Ptch1* were comparable among genotypes (*Smo*, 1.01 ± 0.18 in control, n=8, 1.44 ± 0.57 in ETV1het; DKO, n=8, 1.09 ± 0.52 in TKO, n=8; *Ptch1*, 1.04 ± 0.30 in control, n=8, 1.10 ± 0.34 in ETV1-het; DKO, n=8, 0.80 ± 0.36 in TKO, n=8) (Figure 7B and C). These data suggest that Hedgehog signaling is modulated in the ETV-deficiency-derived cystic kidneys.

**Figure 7.**
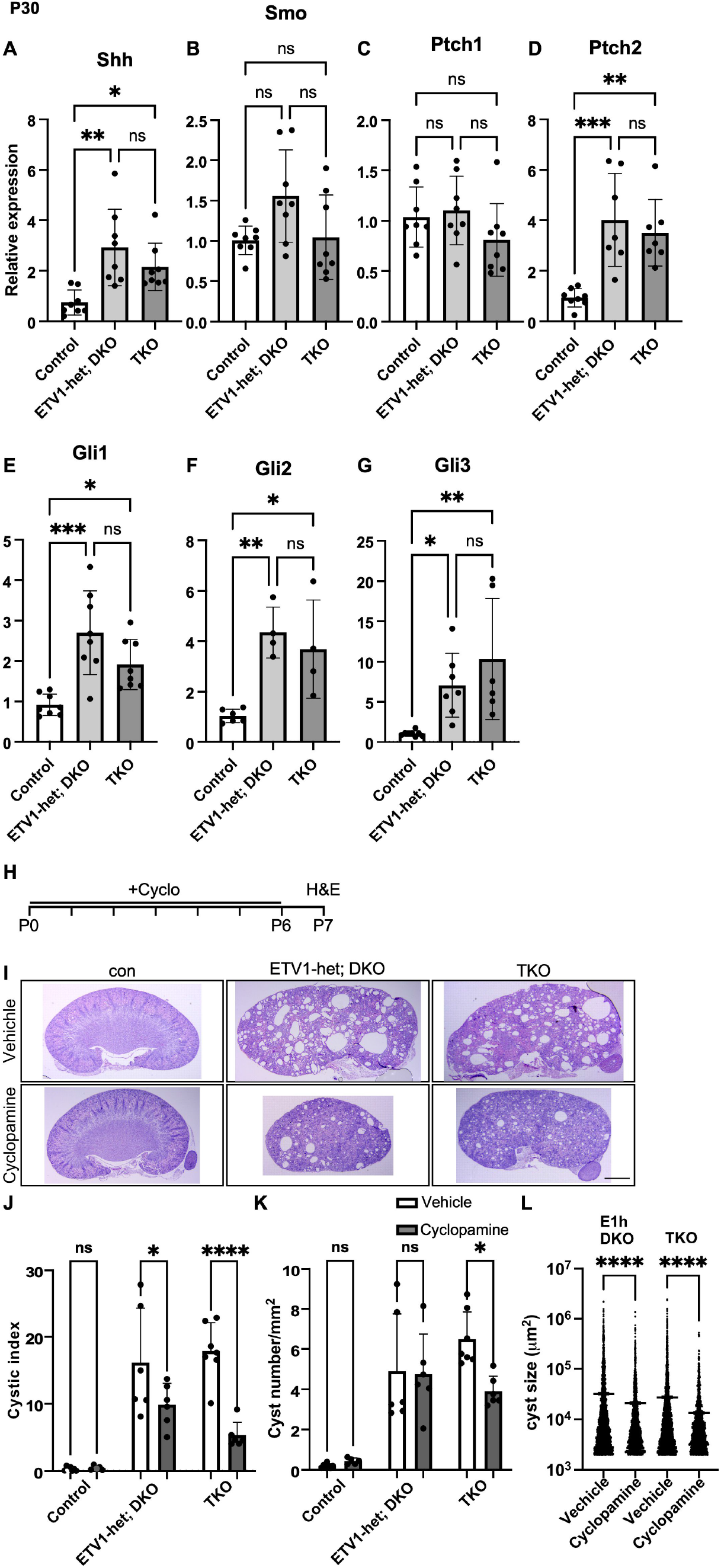
Hedgehog signaling is required for cyst growth caused by loss of ETVs. (A-G) Real-time PCR analyses of Shh (A), Smo (B), Ptch1 (C), Ptch2 (D), Gli1 (E), Gli2 (F), and Gli3 (G) in P30 control, ETV1-het; DKO, and TKO kidneys. (H) Schematic of cyclopamine (3mg/g) treatment. (I) H&E staining of P7 control, ETV1-het; DKO, and TKO treated with/without cyclopamine treatment. (J and K) analyses of cystic index (J) and cyst number (K) P7 control, ETV1-Het; DKO, and TKO treated with/without cyclopamine treatment. *P < 0.05, **P < 0.005, P < 0.0005, ****P < 0.0001. Scale bar, 1mm

Next, we investigated whether decreasing Hedgehog signaling alleviated kidney cyst formation and (or) progression of cyst growth by treating newborn pups with cyclopamine, the inhibitor of Smo, (IP injection, 3mg/kg) for 7 days. H&E staining showed decreased cystic phenotypes in kidneys of ETV1-het; DKO and TKO mice treated with cyclopamine compared to vehicle treated mice (Figure 7I). Further analyses showed that cystic index was significantly decreased by cyclopamine in both ETV1-het; DKO and TKO mice compared to vehicle treated ETV1-het; DKO and TKO mice (0.27 ± 0.30 in vehicle control, n=7, 0.44 ± 0.31 in cyclopamine treated control, n=5, 16.12 ± 8.16 in vehicle ETV1-het; DKO, n=6, 9.85 ± 3.21 in cyclopamine treated ETV1-het; DKO, n=6, P=0.0355, 17.87 ± 4.25 in vehicle treated TKO, n=7, 5.31 ± 1.95 in cyclopamine treated TKO, n=6, P<0.0001) (Figure 7J). Cyst number was significantly decreased in the kidneys of TKO mice treated with cyclopamine compared to vehicle treated TKO mice (0.18 ± 0.11 in vehicle control, n=7, 0.44 ± 0.13 in cyclopamine treated control, n=5, 4.90 ± 2.85 in vehicle ETV1-het; DKO, n=6, 4.75 ± 1.99 in cyclopamine treated ETV1-het; DKO, n=6, 6.48 ± 1.36 in vehicle treated TKO, n=7, 3.90 ± 0.75 in cyclopamine treated TKO, n=6, P=0.0156) (Figure 7K). Cyst size was significantly reduced in cyclopamine treated ETV1-het; DKO and TKO mice compared to vehicle treated mice (64570.18 ± 1749007.29 μm^2^ in vehicle ETV1-het; DKO, n=2923 combined with 7 animals, 42540.83 ± 958161.21 μm^2^ in cyclopamine treated ETV1-het; DKO, n=2018 combined with 6 animals, P<0.0001, 54852.81 ± 1882004.61 μm^2^ in vehicle treated TKO, n=4696 combined with 8 animals, 26847.79 ± 560136.43 μm^2^ in cyclopamine treated TKO, n=1735 combined with 6 animals, P<0.0001) (Figure 7L). These data indicate that inactivation of hedgehog signal reduces cyst progression but has only limited effects on cyst initiation.

## Discussion

In this study, we elucidated a novel function of ETV transcription factors in the cells of nephron compartments. Roles of ETV transcription factors during UB development have been extensively studied. ETV4 and ETV5 were first identified through *ex vivo* screening of Ret/GDNF dependent UB branching morphogenesis [8]. Upon GDNF treatment in the UB explants, ETV5 and ETV5 were induced [8]. Either hypomorphs of ETV4/5 compound mutants or UB specific deletion of ETV4/5 resulted in renal aplasia or hypoplasia due to defects of UB branching [8, 22]. Further studies show that ETV4, expressing the tip of the UB, is responsible to direct UB branching upon GDNF induction [9, 10]. In addition, a case study indicated that homozygous *ETV4* truncation resulted in congenital abnormalities of the kidney and urinary tract [23].

Despite well-characterized roles of ETV transcription factors in the UB, their roles in NPC have been little elucidated. In zebrafish, *Etv5* is dominantly expressed in the multi-ciliated cell domain within the PCT segment and both *Etv4* and *Etv5* are required for multi-ciliated cell identity [11]. In addition, our previous study shows that expression of *Etv4* and *Etv5* are diminished in FGF9/20 double mutant kidneys [12, 13]. In contrast to our expectation, the present study suggests that ETV transcription factors are dispensable for NPC maintenance. Instead, loss of ETV transcription factors leads to generation of multiple cysts during renal development. Lineage and marker studies indicate that renal cystogenesis caused by loss of ETV transcription factors originates from nephron compartments. In addition, ETV4/5 hypomorph kidneys also show renal cysts [8]. These data suggest that ETV transcription factors in the nephron compartment suppress initiation of renal cyst formation.

FGF signaling also regulates renal cystogenesis. Embryonic kidneys of hypomorphs of *Fgf9* and *Fgf2*0 (*Fgf9^-/+^; Fgf20^-/-^*) are cystic in addition to hypoplastic [13]. Nephron compartment specific deletion of *Fgfr2* and its adaptor *Frs2α* results in cystogenesis at the postnatal stage [24, 25]. In addition, these models show that Hedgehog signaling is induced and exhibits cystic kidney disease-like pathologies with acute kidney disease like symptoms, proximal tubule dedifferentiation, and induction of cAMP signaling. Therefore, there are many similarities in renal cystogenesis between loss of FGF signal and ETV transcription factors. Therefore, further investigation is required to understand whether FGF signaling and ETV transcription factors participate the same mechanism regulating renal cystogenesis.

Renal cyst formation is a complex process involving abnormal epithelial cell phenotypes including increased cell proliferation and apoptosis, defects in ciliogenesis, altered matrix adhesion for cell migration, and cell polarity changes [1, 26]. Many of these aspects, including increased cell proliferation and apoptosis, modulation of cilia length, and changes in matrix molecules, are observed in the cysts of ETV mutant kidneys, indicating cysts caused by loss of ETVs share common features of cystogenesis. Therefore, cysts generated through loss of ETV transcription factors could have consequences as bad as other cystic kidney diseases.

Cysts can be classified as simple, malignant, or manifestant based on their size [27]. Although loss ETV4 and ETV5 initiated renal cystogenesis during kidney development, most of the cysts regressed and small number of cysts further progressed (Figure 8). We identified that the regression of initial cysts in the DKO kidneys was due to compensation by ETV1, which was ectopically induced in DKO kidneys. Importantly, cyst number was maintained in the adult when either one copy or two copies of *Etv1* was deleted in DKO kidneys. Either loss of ETV1/5 or ETV1/4 does not result in renal cystogenesis, indicating dispensability of ETV1 on the initiation of cystogenesis. This is one piece of evidence that initiation and promotion of cystogenesis can be separated.

**Figure 8.**
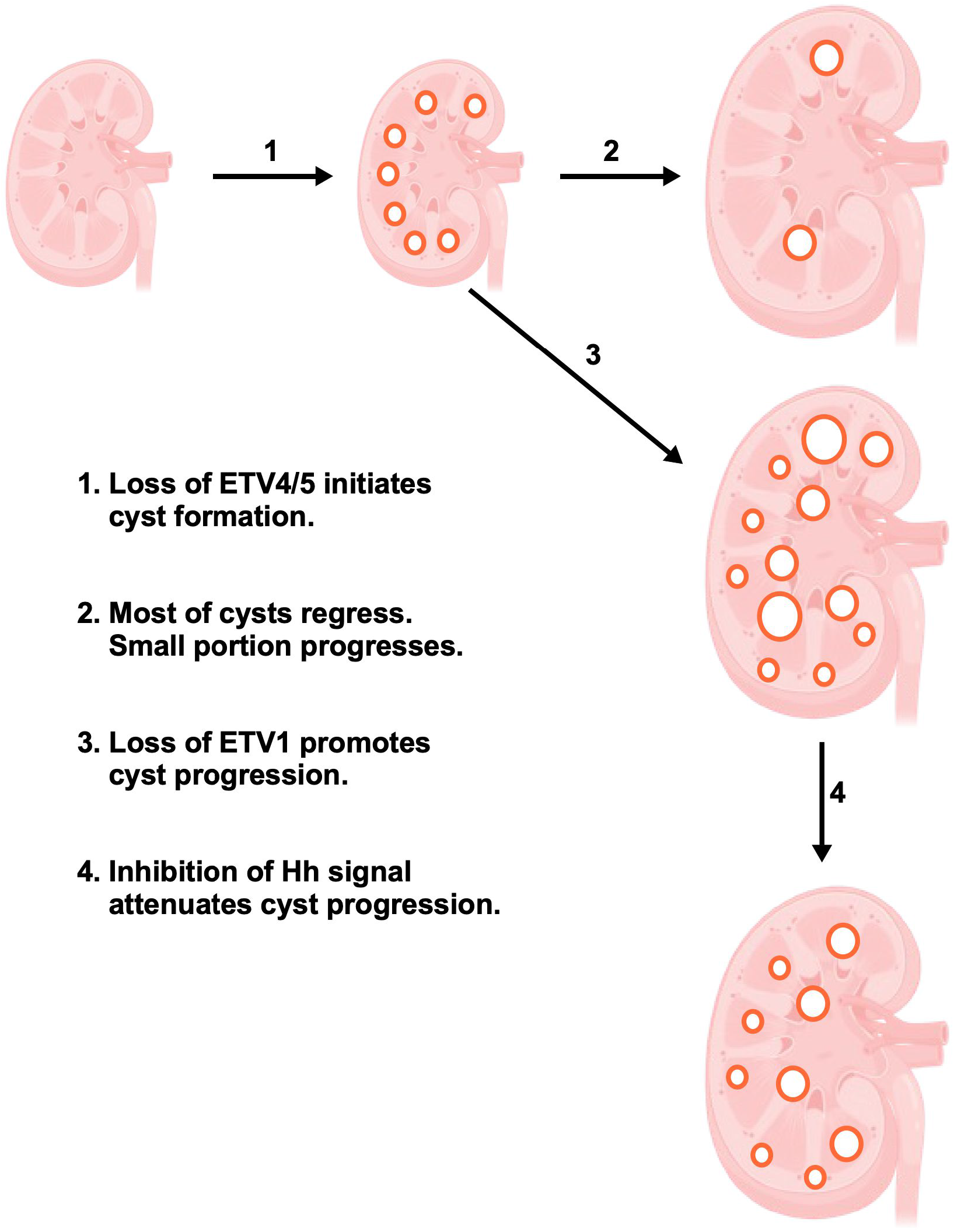
Schematic model of ETV-dependent renal cyst initiation and progression. 1. Renal cyst initiates upon deletion of ETV4 and ETV5 from the nephron compartment. 2. Most of the initial cysts regress throughout the lifetime and only small portion remain to progress. 3. When additional loss of ETV1 occurs, most of the initial cysts start to progress and causes life-threatening cystic disease. 4. Hedgehog upregulation is required for cyst progression and treatment of cyclopamine attenuates cyst progression but not initiation.

Many cystic diseases are caused by mutation of polycystic kidney genes such as Pkd1 and Pkd2. Autosomal dominant polycystic kidney disease (ADPKD) is caused by single allele mutation of *Pkd1* or *Pkd2* [1, 28]. One of characteristics of ADPKD is the focal and sporadic nature of individual cyst formation. There is growing evidence that in addition to genetic mutation of *Pkd1* and *Pkd2*, a second mutation or stimulus is required for pathogenic cyst growth, which refers to ‘two-hit’ model of cystogenesis [29]. This model explains that another somatic mutation of *Pkd1* (or *Pkd2*), in addition to a germline mutation, leads to pathogenic cyst progression. This phenomenon is evident from human cyst samples and genetic mouse model studies [30–33]. The two-hit model expands not only to single gene but also to signaling pathway and modifiers as genomic analyses identified that another gene mutation is common in ADPKD cysts which scores 93% of mutation rate [34]. This information supports that additional modification of gene(s) to decrease the activity of *Pkd1* (or *Pkd2*) is required for pathogenic growth of cysts. This framework fits with our compound loss of ETV transcription factors models and provides evidence that in addition to the commonly known PKD causing genes, other gene families may also contribute to renal cyst initiation and progression.

In addition, we demonstrated that Hedgehog signaling is increased in cystic kidneys, and inhibition of Hedgehog signaling via cyclopamine administration reduced progression but not initiation of cystogenesis. Hedgehog signaling is known to contribute renal cystogenesis. In human ADPKD tissue, hedgehog target gene was activated, and *in vitro* treatment with Hedgehog inhibitors reduced ADPKD cell proliferation [20]. Hedgehog inhibitor treatment of animals with ciliopathy-induced renal cysts (*Arl13b^f/f^; Ksp^Cre^*) reduced cystic burden [21]. In a rat model of Caroli disease presenting with polycystic kidney pathology, treatment with the Hedgehog inhibitor attenuated cystogenesis [35]. This evidence shows that ectopic activation of Hedgehog signaling is one of the players that promote further cyst growth and attenuating Hedgehog signaling may be a viable target to reduce pathogenic cyst burden.

Taken together, we provide the evidence that renal cyst initiation and progression can be molecularly and cellularly distinct and each of steps can be modulated. This information may be expanded to other renal cystic models and be useful to identify molecules to promote/suppress each step and further utilizes as strategies to repress pathogenic cyst growth.

## Materials and Methods

### Mice

This study was carried out in accordance with the recommendations in the Guide for the Care and Use of Laboratory Animals of the National Institutes of Health. The protocol was approved by the University of Nebraska Medical Center Institutional Animal Care and Use Committee (16-005-02-EP) and the University of Mississippi Medical Center Institutional Animal Care and Use Committee (2023-1332). All efforts were made to minimize animal suffering. *Fgf20^Cre/+^*[36] mouse line was reported previously. *Etv1^fl/+^* [37], *Etv4^-/+^* [38], and *Etv5^fl/+^* [39] were gifts from Drs. Silvia Arber from University of Basel, John Hassell from McMaster University, and Xin Sun from University of California San Diego, respectively. *Etv4^-/+^; Etv5^fl/+^ ; Fgf20^Cre/Cre^*males were mated with *Etv4^-/-^*; *Etv5^fl/fl^* females to generate control (*Etv4^-/+^; Etv5^fl/+^ ; Fgf20^Cre/+^*), ETV4-KO (*Etv4^-/-^; Etv5^fl/+^ ; Fgf20^Cre/+^*), ETV5-KO (*Etv4^-+-^; Etv5^fl/fl^ ; Fgf20^Cre/+^*), and DKO (*Etv4^-/-^; Etv5^fl/fl^ ; Fgf20^Cre/+^*) animals. *Etv1^fl/+^; Etv4^-/+^; Etv5^fl/+^ ; Fgf20^Cre/Cre^* males were mated with *Etv1^fl/fl^; Etv4^-/-^*; *Etv5^fl/fl^* females to generate control (*Etv1^fl/+^; Etv4^-/+^; Etv5^fl/+^ ; Fgf20^Cre/+^*), ETV1het; DKO (*Etv1^fl/+^; Etv4^-/-^; Etv5^fl/fl^ ; Fgf20^Cre/+^*), and TKO (*Etv1^fl/fl^; Etv4^-/-^; Etv5^fl/fl^ ; Fgf20^Cre/+^*) animals. Mice were maintained on a 129X1/SvJ;C57B6/J mixed background.

### Histology

For morphological and histological analysis, kidneys from P0, P30, and P180 animals were dissected and images taken using Zeiss Discovery V8 stereo microscopy. Kidneys were then fixed overnight with 4% paraformaldehyde (PFA) and stored in 70% ethanol for paraffin processing. The paraffin processed samples were sectioned and stained with hematoxylin and eosin. For Sirius Red staining, deparaffinized slides were incubated with Picrosirius Red solution for 1hr, and washed twice with acidified water (0.5% acetic acid in water). Stained samples were imaged using a brightfield microscope.

### Immunofluorescence staining

P0, P30, and P180 kidneys were incubated with series of sucrose solutions (10%, 20%, and 30%). Samples were frozen sectioned and stored at -80°C for storage. For antibody staining, sections were washed with PBST (PBS + 0.5% Triton-X 100) for 30 min at RT and incubated with blocking solution (5% donkey serum, in PBST) for 1hr at RT. Sections were incubated with primary antibodies in PBST with 1% donkey serum overnight at 4°C. Sections were washed 3X with PBS and incubated with secondary antibodies for 30min at RT. After washing 3X with PBS, slides were mounted with vectorshield (Vector labs). Images were acquired with Zeiss Axioimage Z2 equipped with ApoTome. Antibodies used were Six2 (Proteintech, 11562-1-AP, 1:500), Aquaporin 1 (Invitrogen, AP5-78806, 1:500), Aquaporin 2 (Invitrogen, PA5-388004, 1:500), NKCC2 (Proteintech, 18970-1-AP 1:500), Ki67 (Proteintech, 27309-1-AP, 1:200), Activated caspase3 (Cell Signaling, 9579S, 1:500), Arl1b3 (Proteintech, 17711-1-AP, 1:500), Foxd1 (Santa Cruz Biotech, sc-47585, 1:200), and CD31 (R&D system, AF3628, 1:500). Biotylated-LTL (Vector labs 1:500), Biotylated-DBA (Vector labs 1:500), and Biotylated-PNA (Vector labs 1:500) were also used. Secondary antibodies conjugated with Alexa488, Alexa555, and Alexa647 (Molecular Probes 1:500) were used.

### Image analysis

Cystic index, cyst number, proliferation index, cell death index, and fibrotic index were measured by Fiji-ImageJ. To measure cystic index and cyst number, cysts and cystic tubules larger than 50 μm in diameter were counted (Park et al., 2009). Cilia length was measured by ACDC software in semiautomatic mode and adjusted manually as recommended [40].

### Florescence In situ hybridization (FISH)

For florescence in situ hybridization, we used probes and reagents purchased from Advanced Cell Diagnostics as recommended by the manufacturer. Probes used were Mm-Etv1, (557891), Mm-Etv4 (458121), and Mm-Etv5 (316961). RNAscope 2.5 HD Detection Reagent – RED (322360) was used.

### Measurement of creatinine and BUN

Blood was collected immediately post-mortem by cardiac puncture, heparinized to prevent coagulation (5 µl of 1000 U/ml heparin per sample; Hospira Inc., Lake Forest, IL) and plasma was collected after centrifuging the blood at 9,000 x G for 5 minutes at 4°C. Plasma creatinine and blood urea nitrogen (BUN) concentrations were determined using QuantiChrom^TM^ Creatinine and Urea Assay Kits, respectively, according to the manufacturer’s directions (BioAssay Systems, Hayward, CA).

### Quantitative real time PCR

P0 and P30 kidneys were dissected out in DEPC-treated cold PBS, then placed in RNA extraction buffer. RNA was extracted using RNeasy mini-RNA extraction kit (Qiagen, 74104) according to the manufacturer’s protocol. Reverse transcription of total RNA was performed using GoScript™ Reverse Transcription System (Promega, A5000) according to the manufacturer’s protocol. Quantitative real time PCR was performed using PowerUp™ SYBR™ Green (Applied Biosytems, A25742) according to the manufacturer’s protocol. Primers are included in supplemental table S1.

### RNA sequencing

Next generation RNA sequencing was performed at University of Nebraska Medical Center Sequencing core. Briefly, synthesis of cDNA from 1–3ng of total RNA was performed with the Clontech SMARTer™ Ultra Low RNA Kit (Clontech Laboratories Inc, Mountain View, CA). Sequencing libraries were prepared using the Illumina Paired End Sample Prep Kit (Illumina Inc, San Diego, CA) according to Illumina’s mRNA-Seq protocol. Each cDNA library was sequenced to generate 75 base pair reads on the Illumina NextSeq500 (Illumina). The sequence reads were aligned to the mouse transcriptome (Ensembl version 102) on the forward strand only using kallisto [41]. These samples have an average alignment rate of 91.5%. Alignments were assembled and annotated using tximport [42]. The database DAVID (the Database of Annotation, Visualization and Integrated Discovery) was used for pathway analysis [43, 44].

### Statistical analysis

Prism (GraphPad) was used to perform a Welch’s t-test or a non-parametric Kruskal-Wallis test with Dunn’s test compensating for multiple comparisons where appropriate, Ordinary one-way ANOVA with Tukey’s multiple comparisons test for multiple comparisons or Unpaired t test where appropriate. A value of P<0.05 was considered statistically significant. For comparative analysis of kidney sizes, kidney perimeters were measured and individual measurements were used for statistical analysis. Three or more animals from at least two independent experiments were examined.

## Acknowledgements

We thank Drs. Silvia Arber, John Hassell, and Xin Sun for providing mice for this study. This work was supported by the NIH P20 GM156708-8333 (S.H).

---

**Supplementary Table1.**
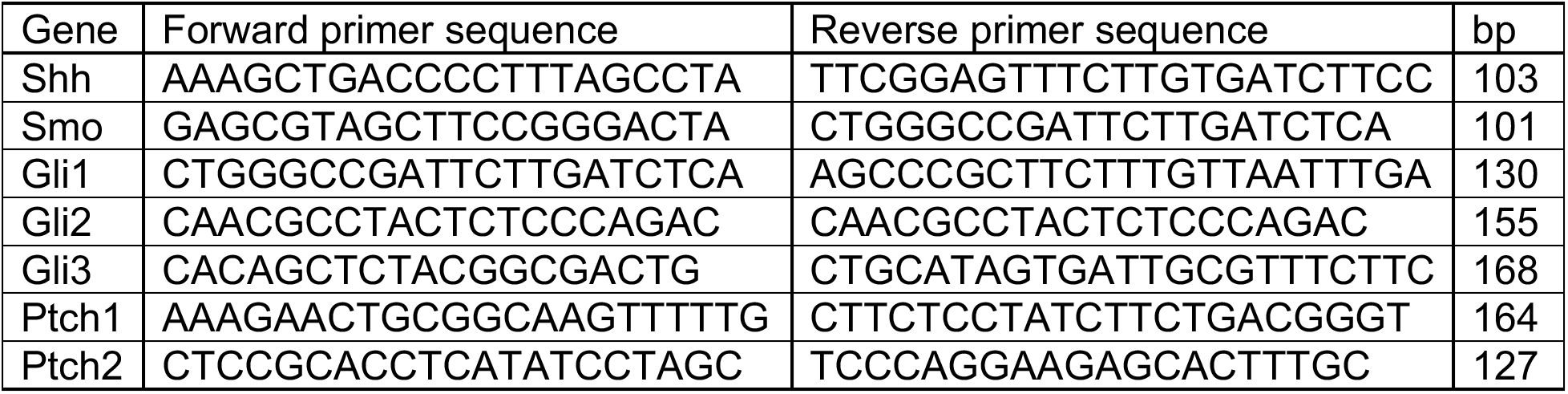
List of primer sets for qRT-PCR.

**SupFigure 1.**
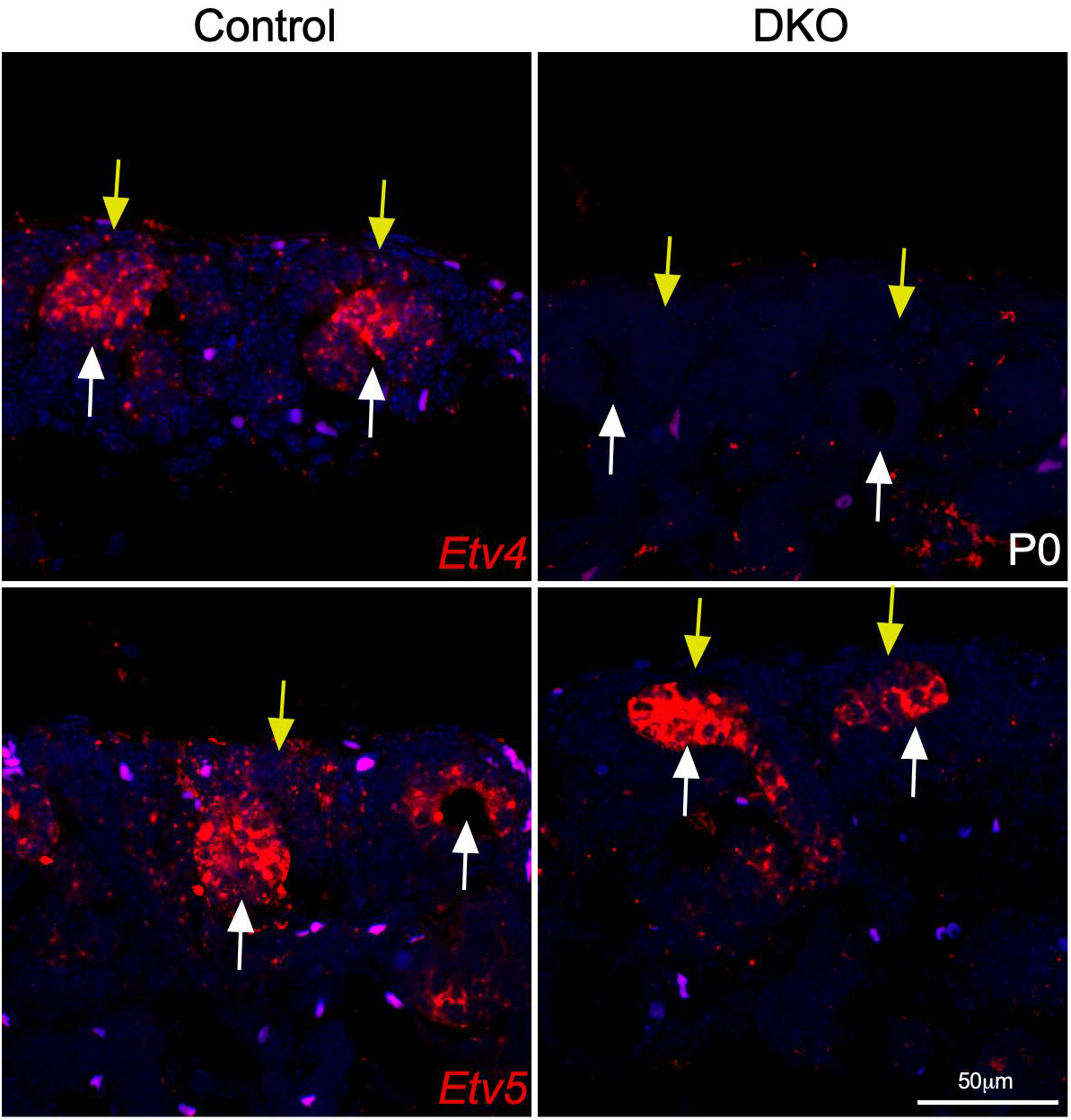
Expression of *Etv4* and *Etv5.* RNA scope of Etv4 and Etv5 at P0 kidney form control and DKO animals. Scale bar 50 µm.

**SupFigure 2.**
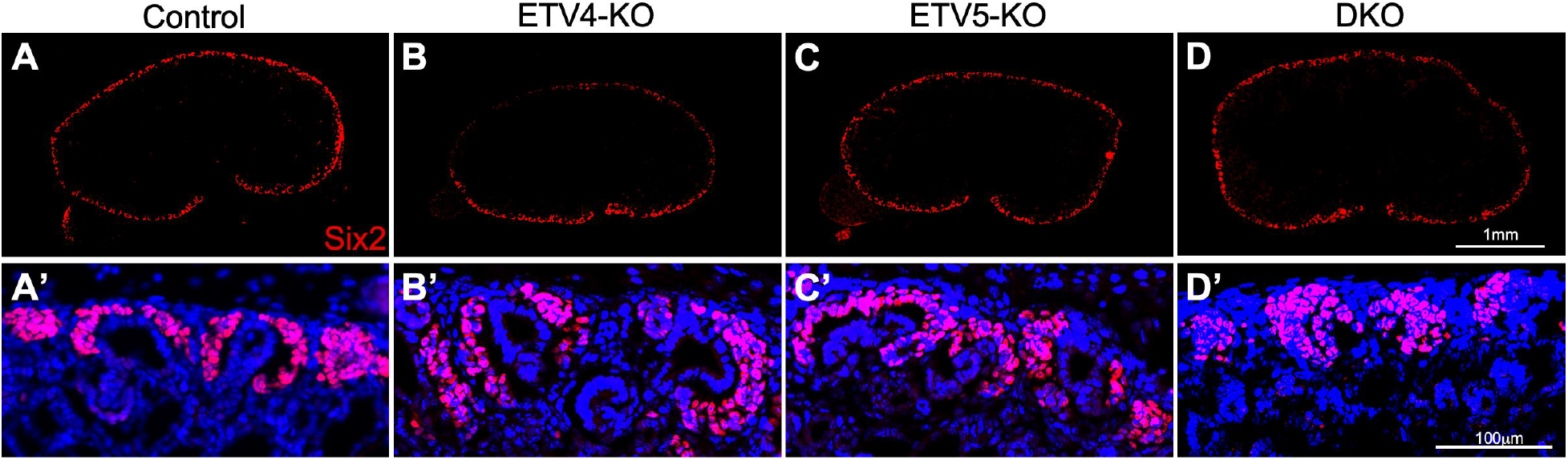
Loss of *Etv4* and *Etv5* in NPC does not influence NPC development. (A-D) Six2 antibody staining showing NPCs in control (A), ETV4-KO (B), ETV5-KO (C), and DKO (D) kidneys showing comparable expression on all genotypes. A’-D’ are inset of A-D. Scale bar A-D, 1mm., A’-D’, 100µm.

**SupFigure 3.**
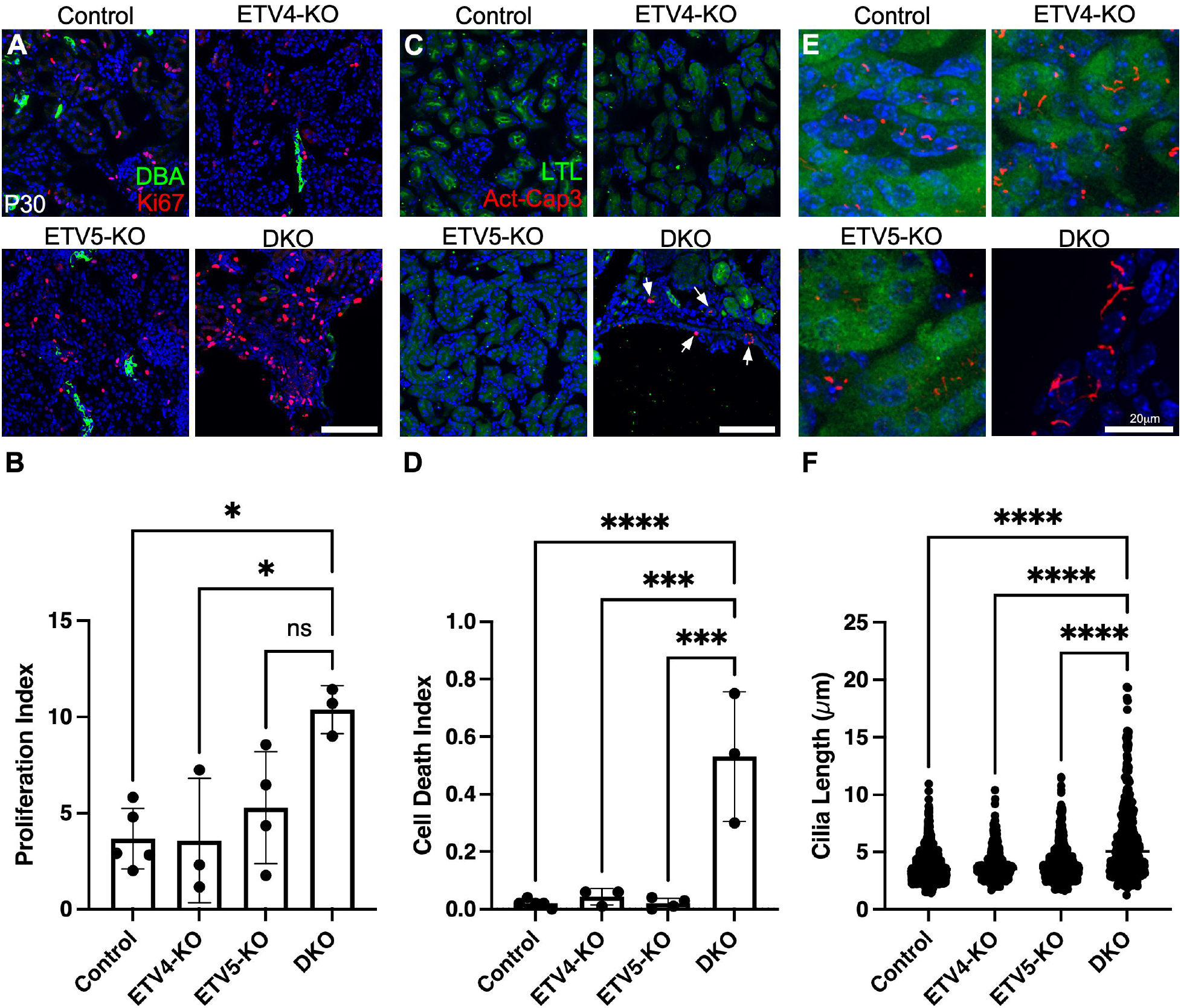
Cystic kidneys increase cell proliferation, cell death, and cilia length at P30. (A-F) Immuno-staining of controls, ETV4-KO, ETV5-KO, and DKO kidneys with LTL and Ki67 for cell proliferation (A), LTL and Activated-Caspase 3 for cell death (Act-Casp3) (C), and LTL and Arl13b (E). Quantification of proliferation index (B), Cell death index (D), and cilia length (F) showing increased in DKOs but no other genotypes. *P < 0.5, ****P < 0.0001. Scale bar, 100µm in A and C, 20µm in E.

**SupFigure 4.**
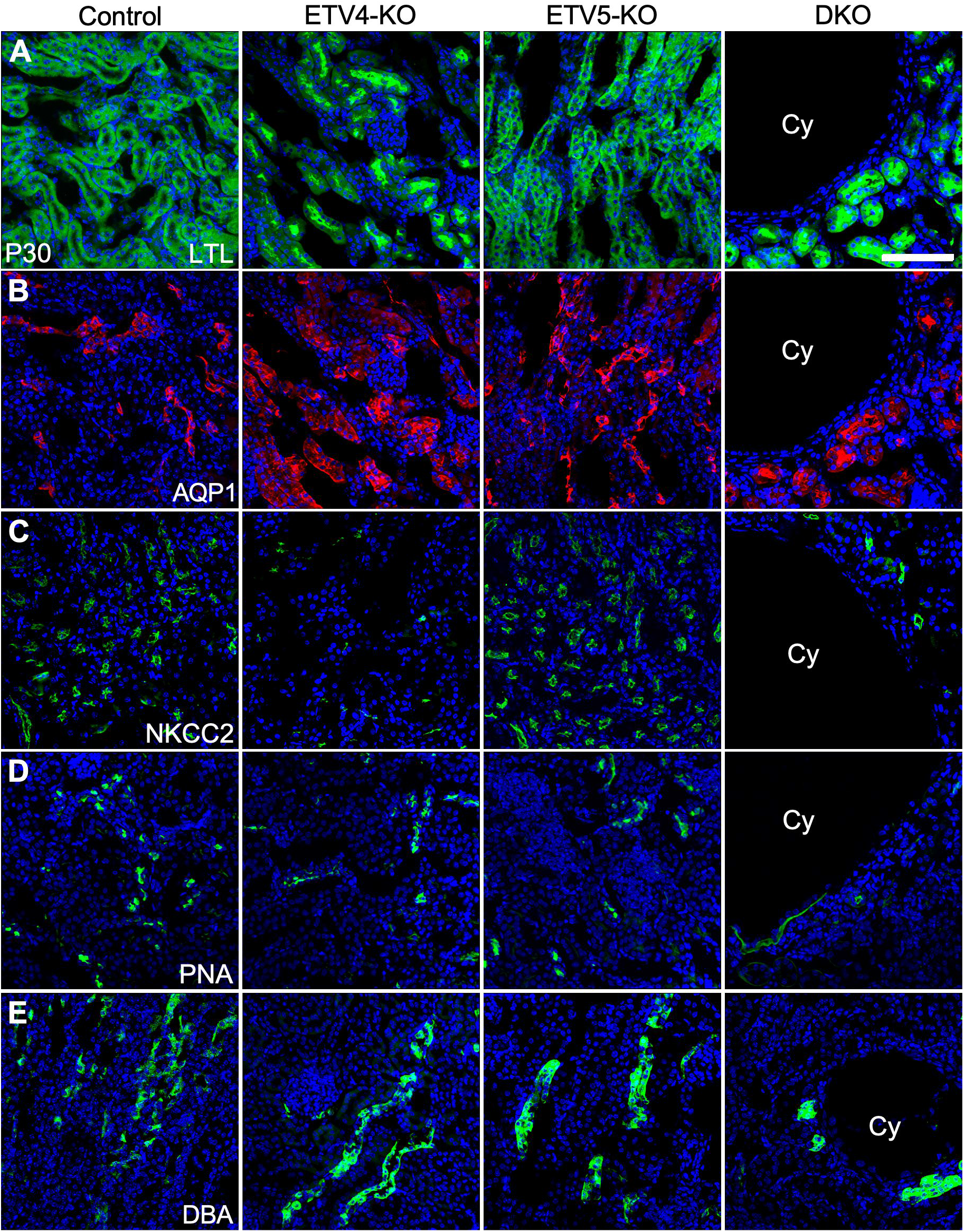
Origin of renal cysts at P30. (A-E) Immuno-staining of markers indicating specific kidney compartments. LTL (A), AQP1 (B), NKCC2 (C), PNA (D), and DBA (E). Cystic cells (Cy) are labelled only with PNA, DCT marker. Scale bar, 1mm in A and B, 100µm in F-J.

**SupFigure 5.**
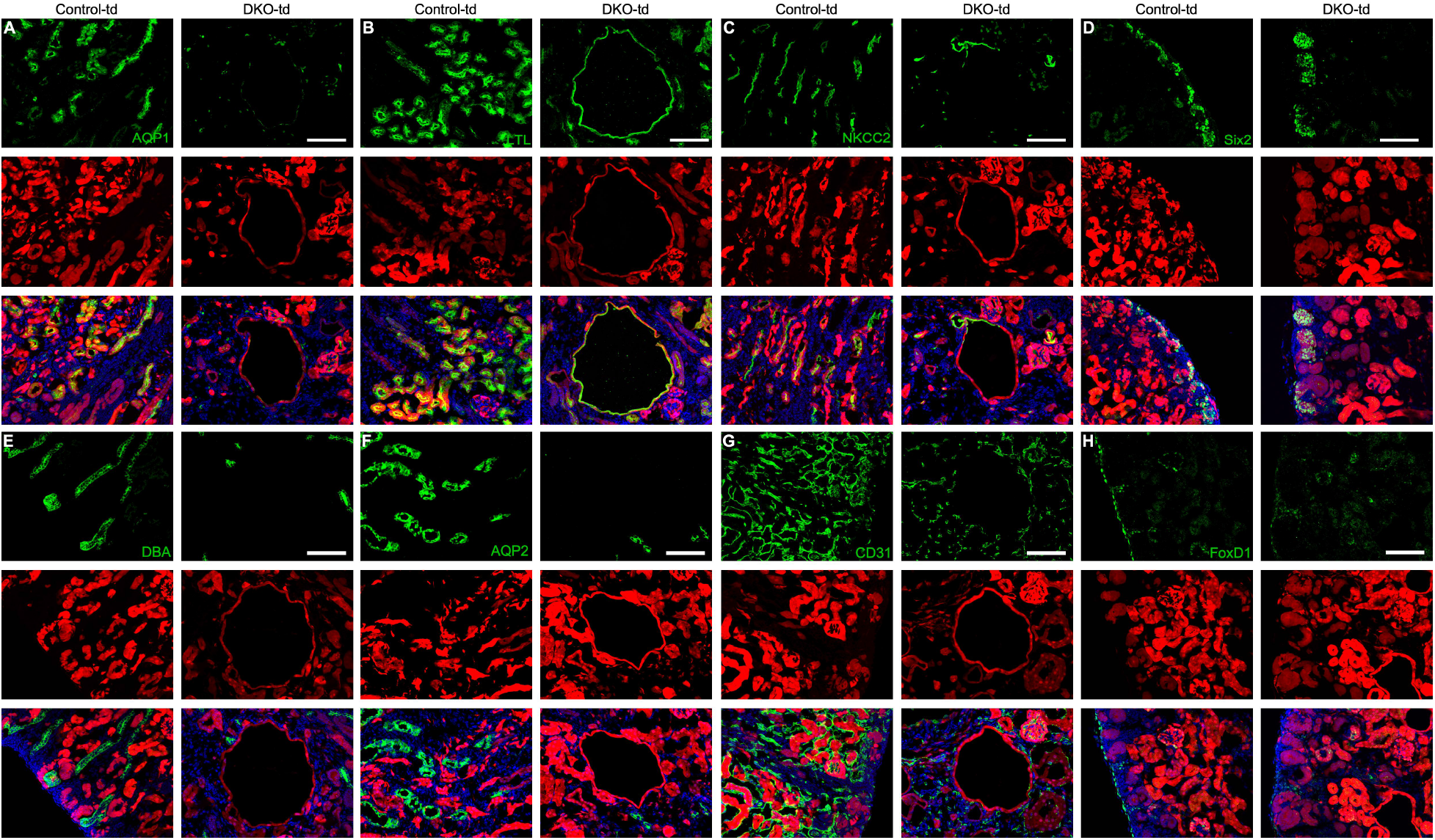
Lineage of renal cysts at P0. (A-H) staining of Aquaporin (A), LTL (B), NKCC2 (C), Six2 (D), DBA (E), Aquaporin 2 (F), CD31 (G), and Foxd1 (H) with tdTomato in P0 control-td (*Etv4^-/+^; Etv5^fl/+^; Fgf20^Cre/+^; Rosa^tdTomato/+^*) and DKO-td (*Etv4^-/-^; Etv5^fl/fl^; Fgf20^Cre/+^; Rosa^tdTomato/+^*) showing NPCs and all nephron compartments are labeled with tdTomato but not ureteric bud cells (E and F), endothelial cells (G), and fibroblast progenitors (H). Scale bar, 1mm.

**SupFigure 6.**
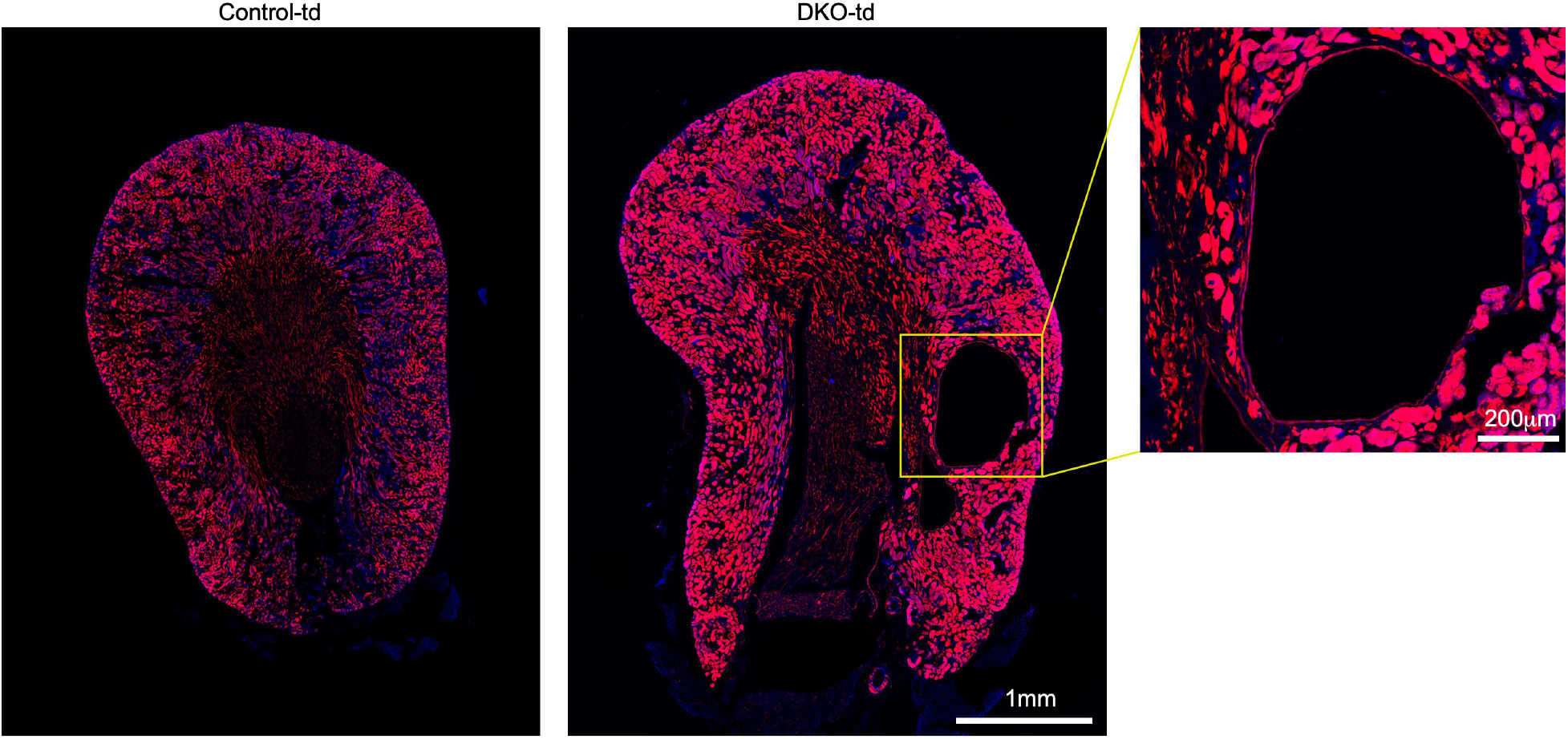
Lineage of renal cysts at P30. tdTomato detection in P30 DKO-td kidney showing the cyst cells are tdTomato positive. Scale bar, 1mm and 200 μm in set.

**SupFigure 7.**
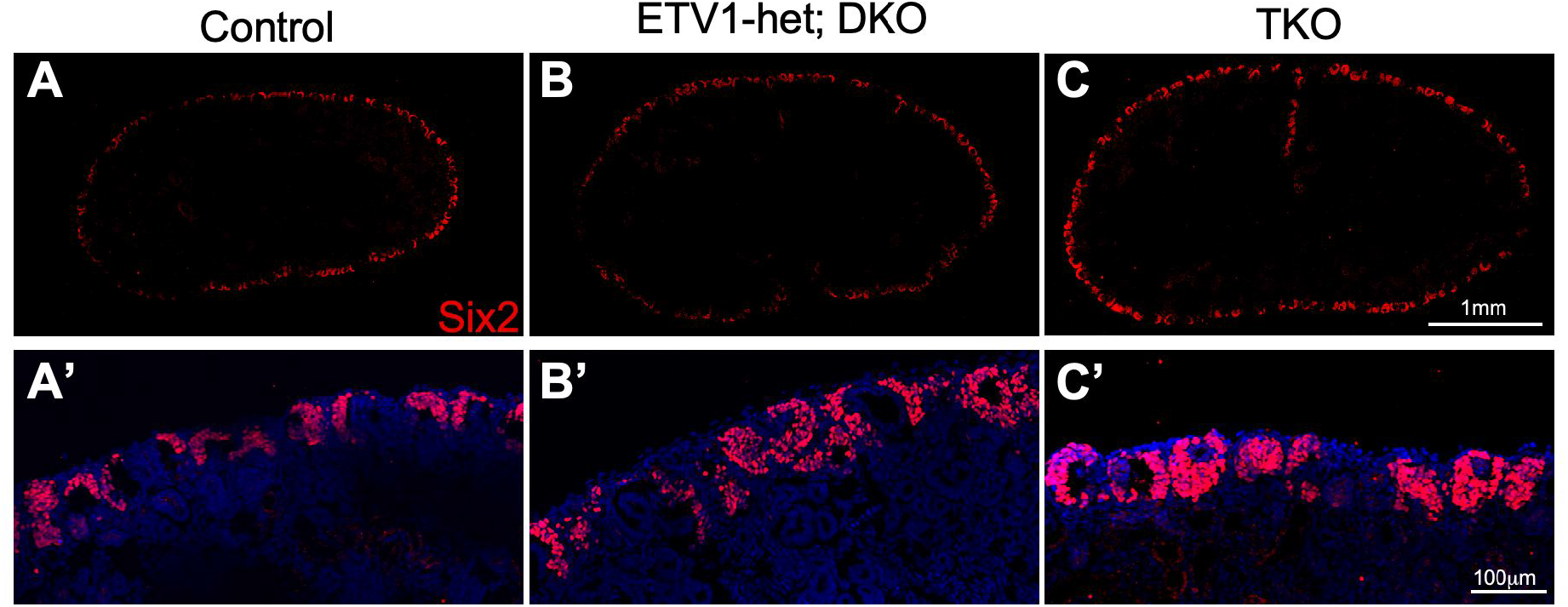
Loss of *Etv1*, *Etv4* and *Etv5* in NPC does not influence NPC development. (A-C) Six2 antibody staining showing NPCs in control (A), ETV1-het; DKO (B), and TKO (C) kidneys showing comparable expression on all genotypes. A’-C’ are inset of A-C. Scale bar A-C, 1mm., A’-C’, 100µm.

**SupFigure 8.**
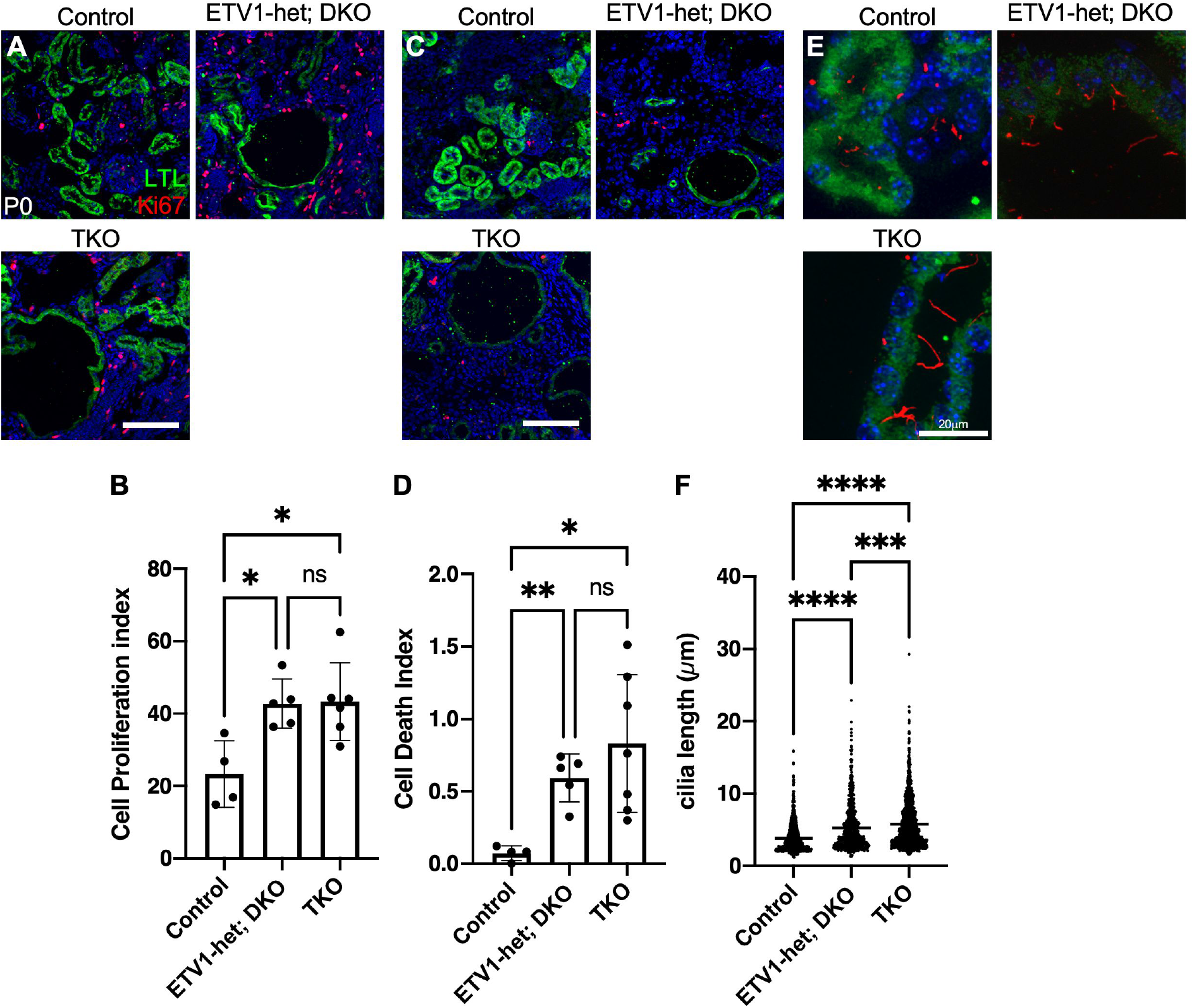
Cystic kidneys increase cell proliferation, cell death, and cilia length at P0 ETV1/4/5 compound kidneys. (A-F) Immuno-staining of controls, ETV1-het; DKO, and TKO kidneys with LTL and Ki67 for cell proliferation (A), LTL and Activated-Caspase 3 for cell death (Act-Casp3) (C), and LTL and Arl13b (E). Quantification of proliferation index (B), Cell death index (D), and cilia length (F) showing increases in ETV1-het; DKO and TKO kidneys compared to control. *P < 0.05, **P < 0.005, ***P < 0.0005, ****P < 0.0001. Scale bar, 100µm in A and C, 20µm in E.

**SupFigure 9.**
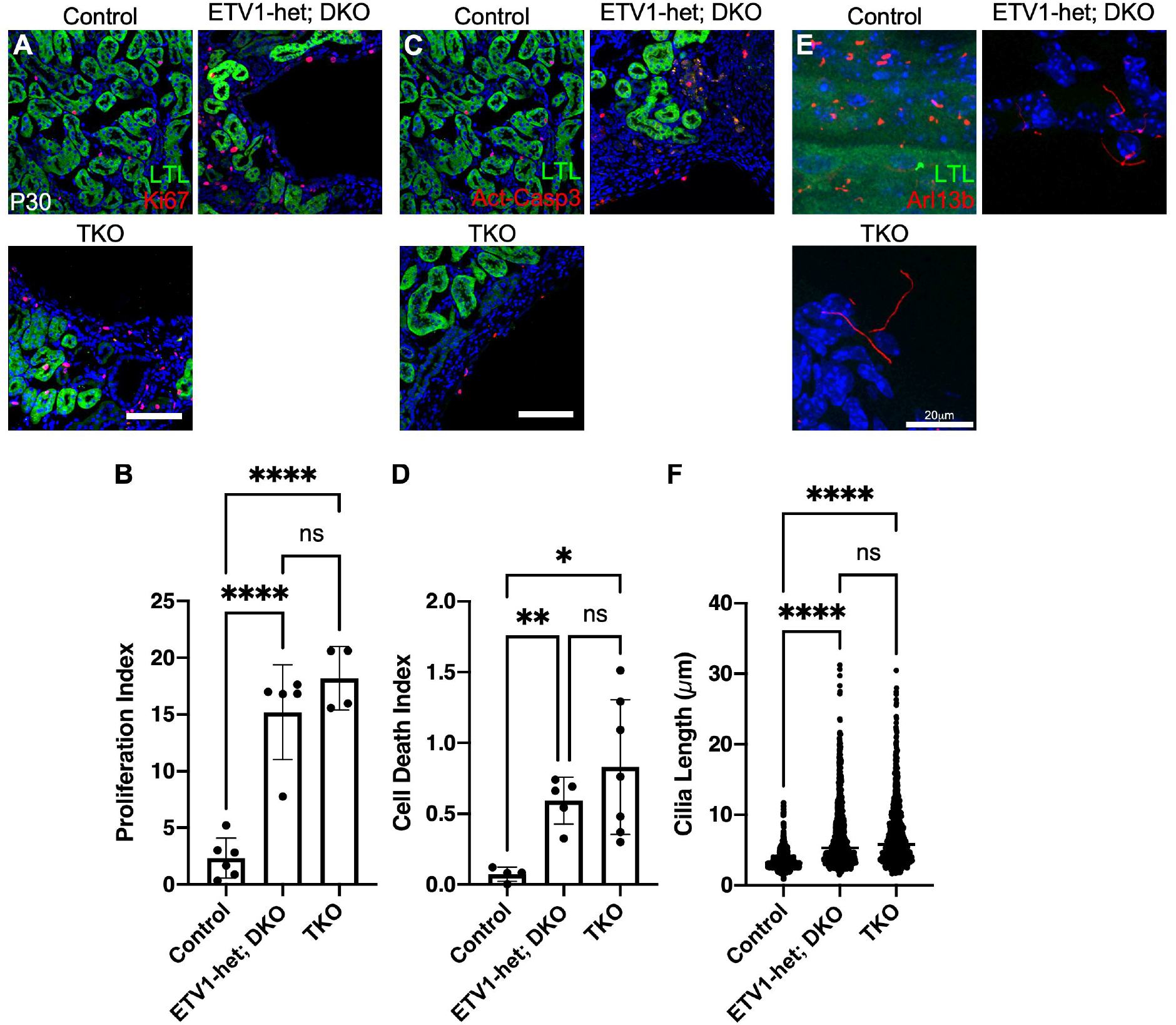
Cystic kidneys increase cell proliferation, cell death, and cilia length at P30 ETV1/4/5 compound kidneys. (A-F) Immuno-staining of controls, ETV1-het; DKO, and TKO kidneys with LTL and Ki67 for cell proliferation (A), LTL and Activated-Caspase 3 for cell death (Act-Casp3) (C), and LTL and Arl13b (E). Quantification of proliferation index (B), Cell death index (D), and cilia length (F) showing increases in ETV1-het; DKO and TKO kidneys compared to control. *P < 0.05, **P < 0.005, ****P < 0.0001. Scale bar, 100µm in A and C, 20µm in E.

**SupFigure 10.**
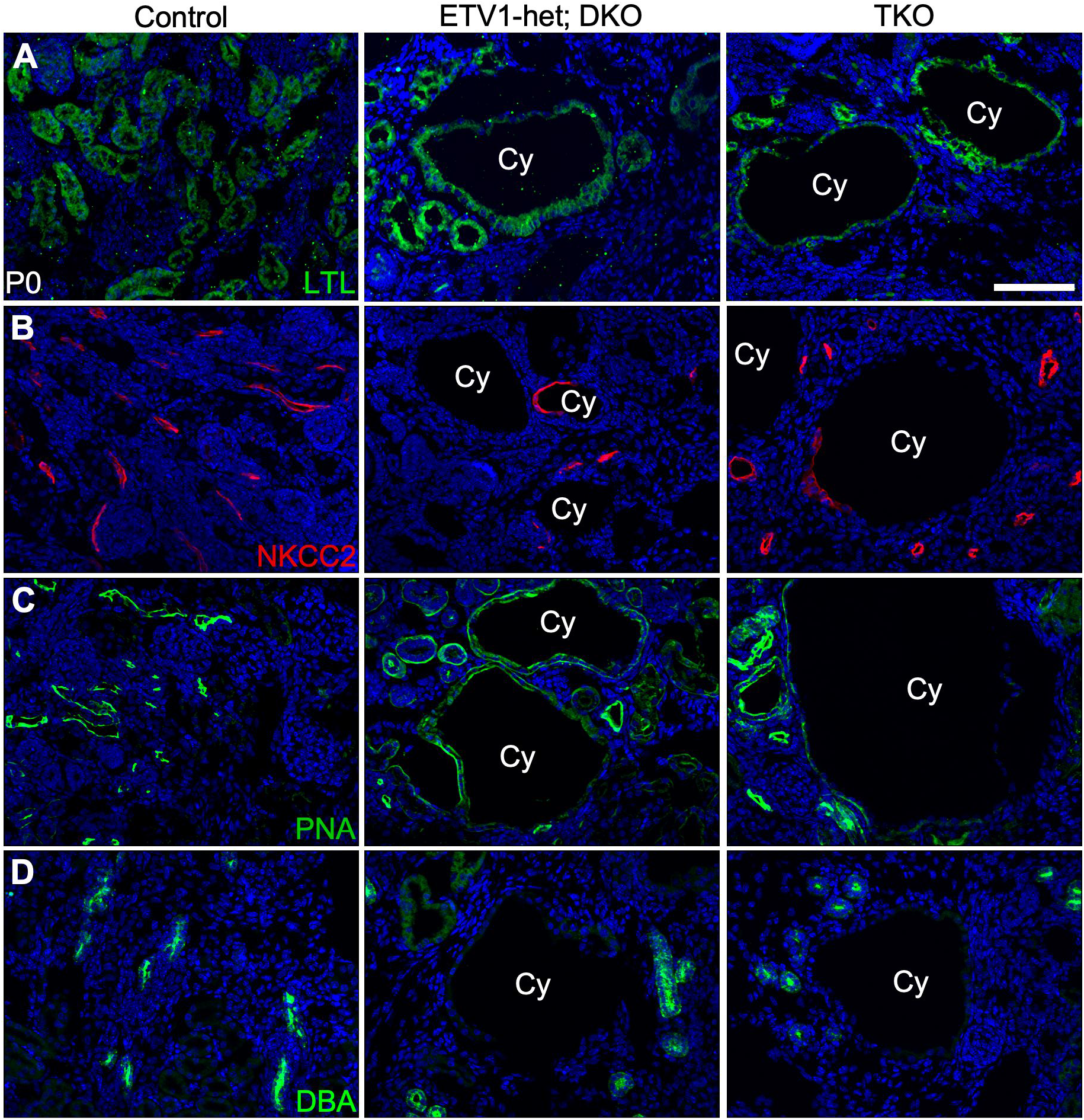
Origin of renal cysts at P0 ETV1/4/5 compound kidneys. (A-D) Immuno-staining of markers indicating specific kidney compartments. LTL (A), NKCC2 (B), PNA (C), and DBA (D). Cystic cells (Cy) are labelled with nephron compartment markers but not UB markers (D). Scale bar, 1mm in A and B, 100µm in F-J.

**SupFigure 11.**
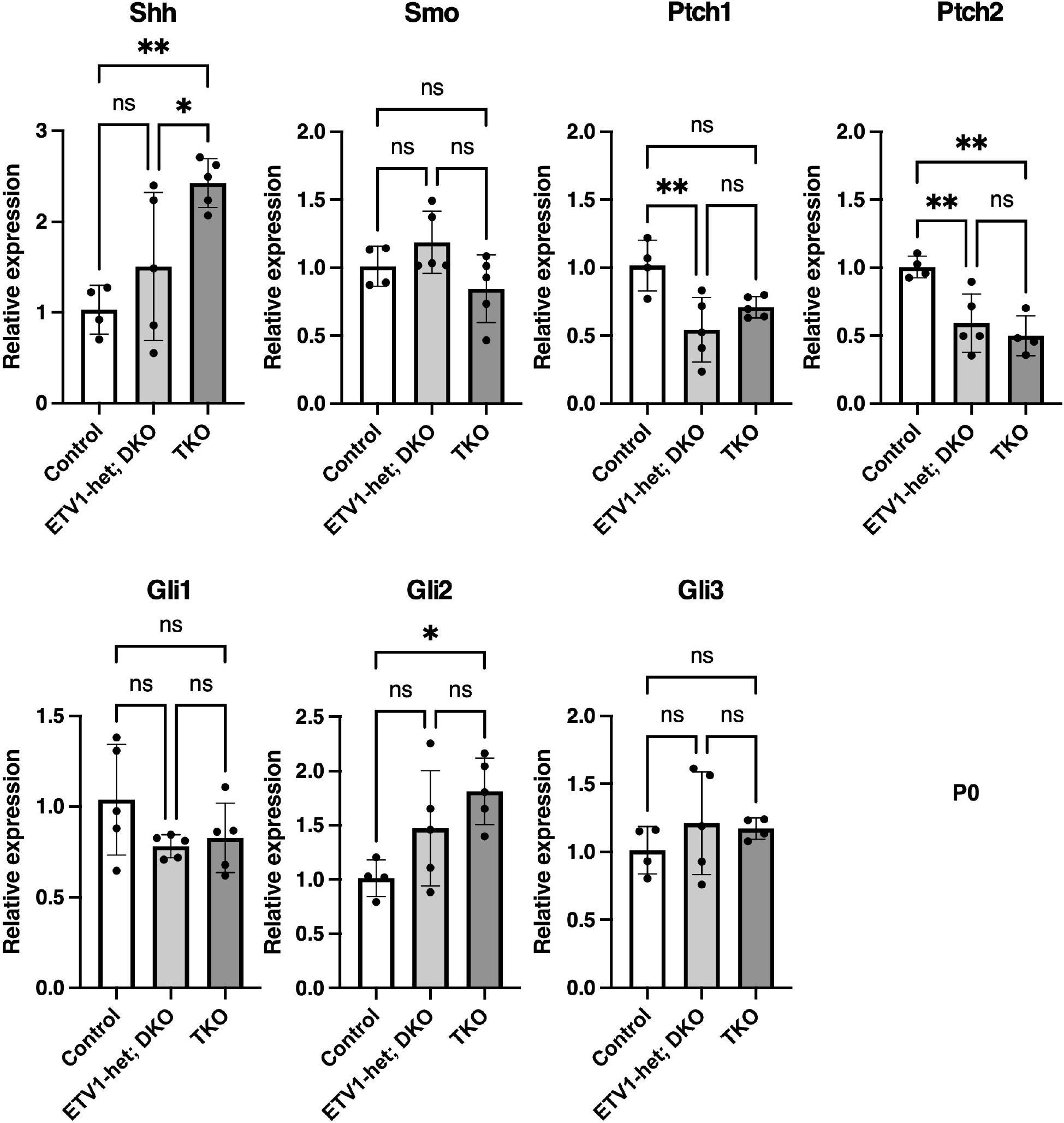
Analysis of hedgehog signal genes from P0 ETV1/4/5 compound kidneys. Real-time PCR analyses of Shh, Smo, Ptch1, Ptch2, Gli1, Gli2, and Gli3 at P0 control, ETV1-het; DKO, and TKO kidneys. *P < 0.05, **P < 0.005.

## Notes

### Competing Interest Statement

The authors have declared no competing interest.

